# Coordinated interactions between endothelial cells and macrophages in the islet microenvironment promote β cell regeneration

**DOI:** 10.1101/2020.08.28.265728

**Authors:** Diane C. Saunders, Kristie I. Aamodt, Tiffany M. Richardson, Alec Hopkirk, Radhika Aramandla, Greg Poffenberger, Regina Jenkins, David K. Flaherty, Nripesh Prasad, Sean E. Levy, Alvin C. Powers, Marcela Brissova

## Abstract

Endogenous β cell regeneration could alleviate diabetes, but proliferative stimuli within the islet microenvironment are incompletely understood. We previously found that β cell recovery following hypervascularization-induced β cell loss involves interactions with endothelial cells (ECs) and macrophages (MΦs). Here we show that proliferative ECs modulate MΦ infiltration and phenotype during β cell loss, and recruited MΦs are essential for β cell recovery. Furthermore, VEGFR2 inactivation in quiescent ECs accelerates islet vascular regression during β cell recovery and leads to increased β cell proliferation without changes in MΦ phenotype or number. Transcriptome analysis of β cells, ECs, and MΦs reveals that β cell proliferation coincides with elevated expression of extracellular matrix remodeling molecules and growth factors likely driving activation of proliferative signaling pathways in β cells. Collectively, these findings suggest a new β cell regeneration paradigm whereby coordinated interactions between intra-islet MΦs, ECs, and extracellular matrix mediate β cell self-renewal.

## INTRODUCTION

While investigating how vascular endothelial growth factor-A (VEGF-A) regulates islet vascularization, our group previously characterized a mouse model in which signals from the local microenvironment stimulate β cell self-renewal^1^. In this model, transiently increasing VEGF-A production in β cells (βVEGF-A) induces endothelial cell (EC) expansion and hypervascularization that causes β cell loss; remarkably, islet morphology, capillary network, β cell mass, and function normalize 6 weeks after withdrawal (WD) of the VEGF-A stimulus. This regenerative response is a result of a transient but robust burst in β cell proliferation, which is dependent on VEGF-A-mediated recruitment of macrophages (MΦs). These recruited cells express markers of both pro-inflammatory (M1) and restorative (M2) activation, suggesting a unique regenerative phenotype^1–5^. Further investigation into the role of various microenvironmental components on β cell proliferation in this model is needed given that this regenerative microenvironment promotes proliferation of human in addition to mouse β cells^1^.

MΦs are often perceived as damaging to islets due to their role in β cell loss during diabetes^6–8^, but it is becoming increasingly appreciated that tissue-resident MΦs play important roles in immune surveillance and tissue homeostasis and function in the islet. Mice with compromised MΦ populations during pancreatic development exhibit reduced β cell proliferation, β cell mass, and impaired islet morphogenesis^9,10^. Furthermore, recent studies have supported the notion that MΦs contribute to β cell regeneration after several types of injury, including surgically induced pancreatitis^11^ and diphtheria toxin (DT)-mediated apoptosis^12^. This recent work highlights the importance of understanding how MΦs contribute to β cell proliferation and further defining MΦ phenotype and function in the βVEGF-A model.

In addition to MΦs, ECs are known to participate in tissue repair via activation of the VEGF-A–VEGFR2 pathway, which mediates angiocrine factor production and promotes local cell renewal and regeneration^13–16^. There is a precedent for ECs facilitating tissue repair by influencing MΦ activation toward a restorative, M2-like phenotype^4^. Because vasculature is essential for normal islet function, understanding signals that govern EC homeostasis and the effects of ECs on neighboring cell populations is crucial for maintaining and restoring islet health. Signaling between ECs and the pancreatic epithelium is critical for establishing islet vasculature and β cell mass during development, and in mature islets ongoing signaling between endocrine and ECs is required to maintain the capillary network through which endocrine cells receive adequate nutrition and oxygen and can rapidly sense and secrete hormones^17–19^. The vascular basement membrane is also the primary component of the intra-islet extracellular matrix (ECM) and acts as a reservoir for growth factors and other signaling molecules important for β cell differentiation, function, and proliferation^20–23^. In the βVEGF-A model, ECs may affect the regenerative process indirectly by promoting MΦ recruitment and activation, altering ECM composition and signaling, and/or by directly influencing β cell proliferation.

Here we deconstructed the complex *in vivo* islet microenvironment in the βVEGF-A model and show that β cell self-renewal is mediated by coordinated interactions between recruited MΦs, intra-islet ECs, and the ECM (**Figures 1, S1**). To isolate the roles of MΦs and ECs in this model we removed MΦs from the islet microenvironment (**Figures 1, S1**; experimental scheme **A**) and inactivated VEGFR2 signaling in ECs to discern the effects of proliferative or quiescent ECs on β cell proliferation (**Figures 1, S1**; experimental schemes **B** and **C**). Since the βVEGF-A islet microenvironment is a complex *in vivo* system involving dynamic changes in islet cell composition, we also identified regenerative signals by performing transcriptome analysis (**Figures 1, S1**; experimental scheme **D**) of purified islet cell populations including β cells, ECs, and MΦs over the course of β cell loss and recovery. Based on previous work^1,16^ we predicted that either MΦ depletion (clodronate) or loss of VEGFR2 signaling in ECs would perturb MΦ recruitment and polarization and impair β cell regeneration. Indeed, we found that MΦ depletion suppressed the transient burst in β cell proliferation leading to reduced recovery of β cell mass.

**Figure 1.**
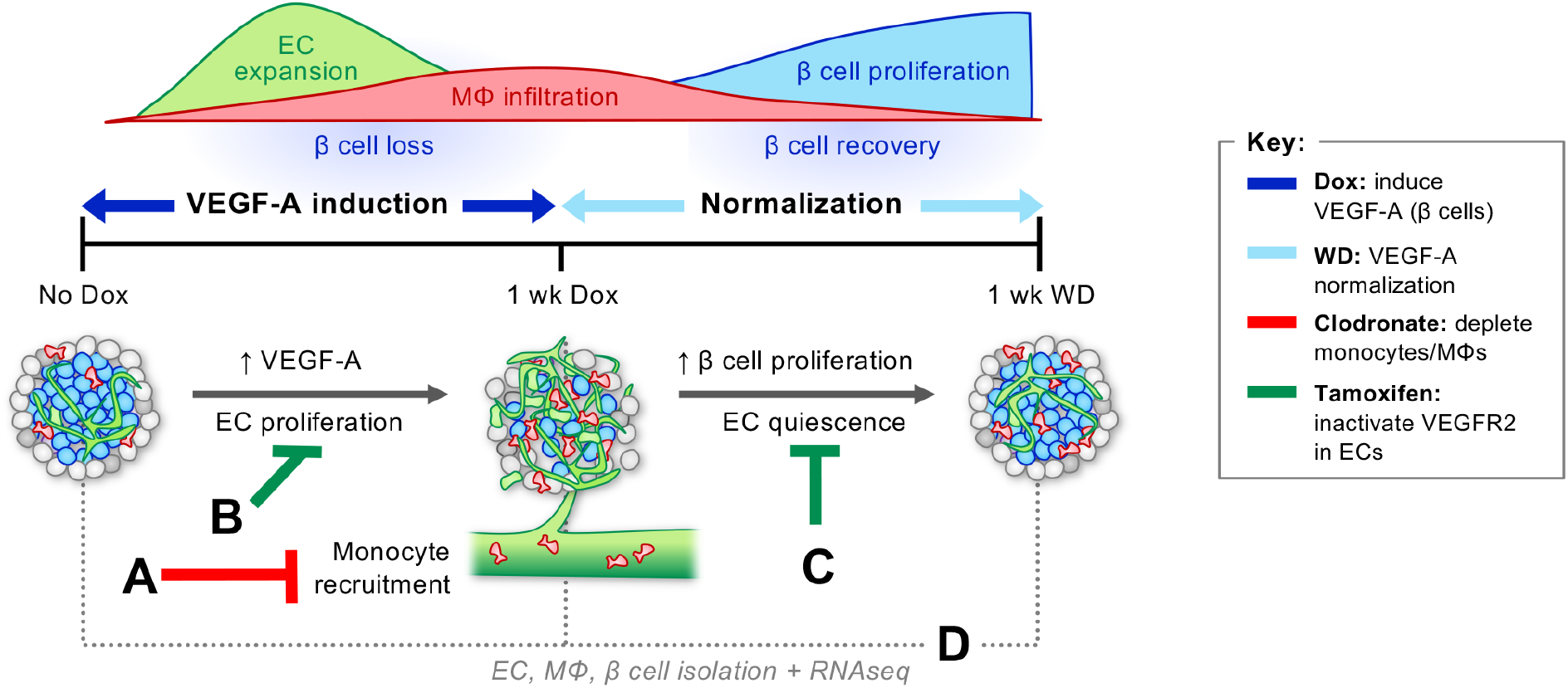
The RIP-rtTA; Tet-O-VEGF-A (βVEGF-A) model of β cell regeneration permits specific modulation of the islet microenvironment. Induction of VEGF-A overexpression in β cells with doxycycline (Dox) causes rapid endothelial cell (EC) expansion, β cell death, and recruitment of circulating monocytes increasing the number of intra-islet macrophages (MΦs). Once the Dox stimulus is removed, VEGF-A levels normalize and β cells undergo self-renewal^1^. **(A)** To establish the role of MΦs in β cell proliferation, monocytes and MΦs were depleted with clodronate liposomes. **(B)** To determine whether proliferative intra-islet ECs were required for β cell loss, MΦ recruitment, and MΦ phenotype activation, an additional genetic construct was introduced to knock down key signaling receptor VEGFR2 in ECs prior to VEGF-A induction. **(C)** To determine if VEGFR2 signaling in quiescent ECs contributes to β cell proliferation, VEGFR2 was inactivated in ECs during β cell recovery. **(D)** To identify cell-specific transcriptome changes during β cell loss and recovery, populations of ECs, MΦs, and β cells were isolated prior to and during VEGF-A induction, and after VEGF normalization. For additional details, including mouse models utilized, see **Figure S1**.

In addition, VEGFR2-mediated signaling in intra-islet ECs was necessary for maximal MΦ recruitment and M2-like phenotype activation. Surprisingly, VEGFR2 ablation in quiescent ECs during the period of β cell recovery accelerated islet vascular regression leading to increased β cell proliferation while MΦ phenotype and number was unchanged. Transcriptome analysis during β cell death and recovery revealed intricate changes in expression of growth factors, integrins, and matrix remodeling enzymes in all three cell types, suggesting that ECM remodeling and activation of ECM-associated molecules within the islet microenvironment play critical roles in β cell self-renewal. Taken together, our results indicate that both MΦs and intra-islet ECs provide crucial microenvironmental cues to cooperatively promote β cell regeneration.

## RESULTS

We previously developed the βVEGF-A model, which allows for Dox-induced, β cell-specific overexpression of VEGF-A causing rapid intra-islet EC expansion, widespread β cell loss, and MΦ recruitment to islets from a pool of circulating monocytes^1^. When Dox is removed and VEGF-A levels normalize, ECs return to baseline levels, but MΦs persist in the islet microenvironment, and β cells undergo a transient but robust proliferation that leads to β cell mass restoration (**Figure 1**).

### Chemical depletion of macrophages in βVEGF-A islets inhibits β cell proliferation

To dissect pathways and signaling molecules in the islet microenvironment required for β cell recovery, we first employed clodronate-mediated MΦ depletion which impacts both infiltrating and islet resident MΦs. Clodronate is an ATP/ADP translocase inhibitor, and when packaged into liposomes it is selectively taken up by MΦs due to their phagocytic properties and causes apoptosis. We treated βVEGF-A mice with either control or clodronate liposomes starting one day before VEGF-A induction and continuing for one week after VEGF-A normalization (**Figure 2A**). Compared to control, clodronate treatment reduced circulating CD11b^+^ Ly6G^−^ monocytes by 50% within 24 hours (**Figure 2B,***No Dox*, and **Figure S2**) and reduced the MΦ population in islets by 94% one week after VEGF-A induction (**Figure 2C** and **2D**, *1wk Dox*). MΦ depletion was maintained during VEGF-A normalization, with 86% fewer MΦ in islets from clodronate-treated βVEGF-A mice one week after Dox withdrawal (**Figure 2C** and **2D**, *1wk WD*). VEGF-A induction in clodronate-treated mice led to increased EC area (**Figures 2C** and **2E**) and β cell loss (**Figures 2B** and **2F**) comparable to βVEGF-A mice treated with control liposomes, thus demonstrating that clodronate treatment and MΦ depletion did not influence intra-islet EC expansion and hypervascularization-induced β cell loss. In contrast, MΦ depletion significantly impaired β cell proliferation during the recovery period (8.5 vs. 2.1%, p<0.001; **Figure 2G**) and resulted in reduced β cell area compared to controls after 6 weeks of VEGF-A normalization (**Figure 2F**), demonstrating that MΦs are required for the regenerative response in βVEGF-A islets. Interestingly, β cell area is slightly but significantly increased one week after Dox withdrawal (**Figure 2F**, *1wk WD*) in clodronate-treated βVEGF-A mice before ultimately the impaired β proliferation leads to reduced β cell area 6 weeks after VEGF-A normalization. This finding suggests that MΦs play an important role in islet remodeling and composition in βVEGF-A mice separate from their effect on β cell proliferation, most likely through their function as phagocytes.

**Figure 2.**
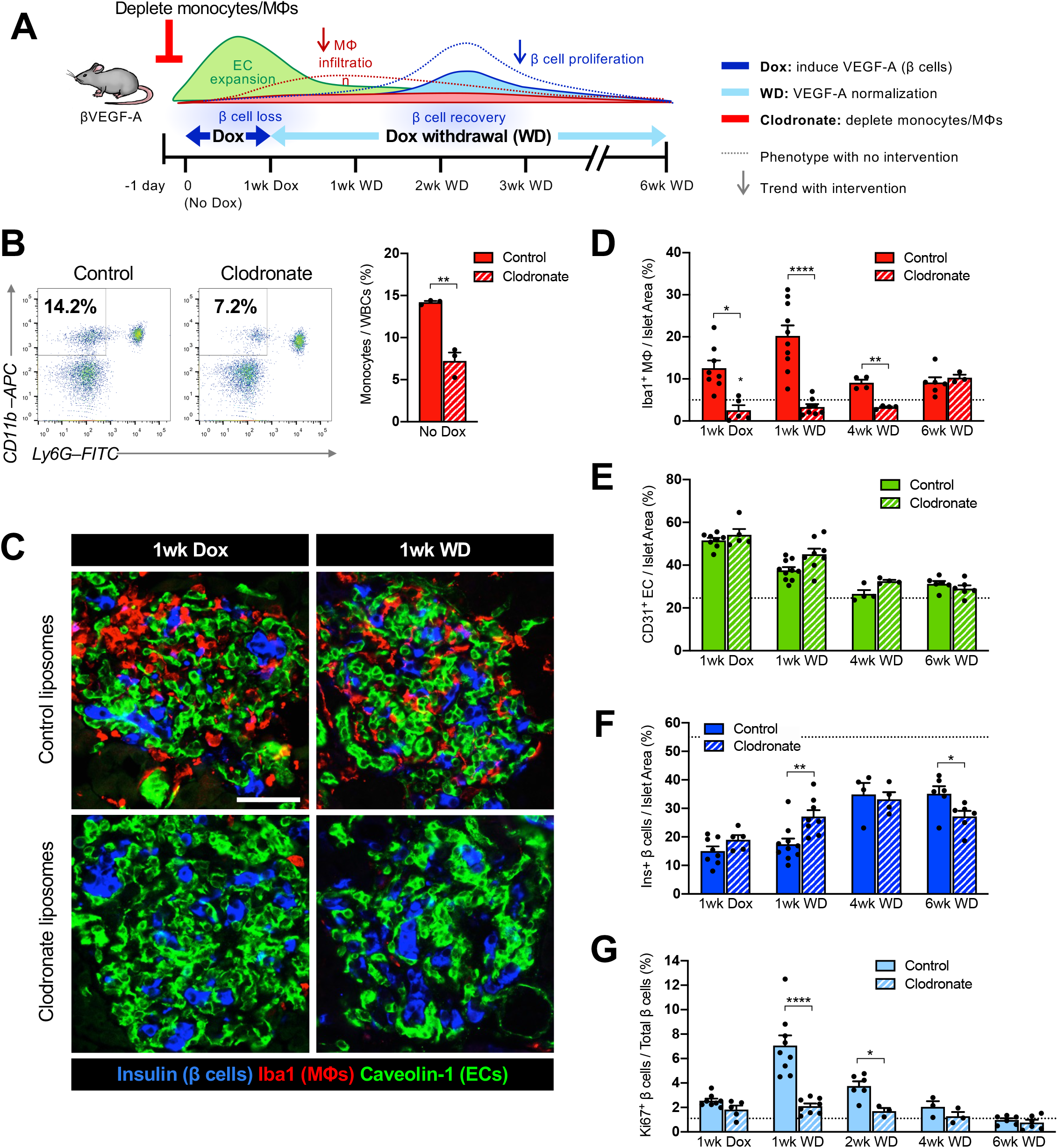
Macrophages are required for β cell proliferation in βVEGF-A mice. **(A)** To deplete macrophages (MΦs) during VEGF-A induction and normalization, βVEGF-A mice were treated with clodronate or control liposomes (150-200 μl i.v.) every other day, beginning one day before Dox treatment and continuing 1 week after Dox withdrawal (1wk WD). **(B)** Representative flow cytometry plots showing circulating monocytes (CD11b^+^ Ly6G^−^) of control and clodronate-treated βVEGF-A mice 24 hours after single injection (No Dox). Approximately 10,000 white blood cells (WBCs) were analyzed (monocyte fraction reported as mean±SEM) per each animal. **(C)** Islet architecture displayed by labeling for β cells (Insulin; blue), endothelial cells (Caveolin-1; green), and MΦs (Iba1; red) during VEGF-A induction (1wk Dox) and normalization (1wk WD). Scale bar, 50 μm. **(D)** Quantification (mean±SEM) of islet MΦ area by immunohistochemistry (5.1±0.7 × 10^5^ μm^2^ total islet area analyzed per animal). **(E–F)** Quantification (mean±SEM) of endothelial cell (EC) area **(E)** and β cell area **(F)** in βVEGF-A mice treated with control or clodronate liposomes during VEGF-A induction and normalization. **(G)** Rate of β cell proliferation (mean±SEM; 1,138±87 β cells counted per animal) during VEGF-A normalization (1wk WD and 2wk WD) in control βVEGF-A mice was significantly reduced in clodronate-treated mice. In panels **(B–G)** each closed circle represents one animal; asterisks indicate unpaired two-tailed t-tests of control vs. clodronate groups; *, p<0.05; **, p<0.01; ****, p<0.0001. Dashed lines in **D–G** depict average values in βVEGF-A mice at baseline (No Dox).

### Proliferative ECs are required for MΦ polarization and maximal MΦ recruitment

To investigate the contribution of ECs to β cell loss and recovery in the βVEGF-A model, we first created a mouse line in which VEGFR2, which is enriched in ECs of islet capillaries and the main transducer of VEGF-A signal in islets, is inactivated by tamoxifen (Tm)-inducible Cre-mediated excision in ECs. We initially tested the EC-SCL-CreER^T^ transgene^24^, which contains a 5’ endothelial enhancer for the stem cell leukemia (SCL) transcription factor, and although Cre activity was confirmed in islet ECs using the *Gt(ROSA)26Sor^tm1Sor^* reporter strain^25^, VEGFR2 expression was unchanged in EC-SCL-CreER^T^; VEGFR2^fl/fl^ mice (data not shown). Fortunately, were able to obtain the Cad5-CreER^T2^ line^26^ and achieved efficient EC-specific VEGFR2 knockdown in Cad5-CreER^T2^; VEGFR2^fl/fl^ (VEGFR2^iΔEC^) mice. Pancreata harvested after 3 doses of Tm were evaluated to confirm VEGFR2 ablation (**Figures S3A** and **S3B**) without significant changes in islet capillary density or size (**Figures S3C** and **S3D**) or basal β cell proliferation (**Figure S3E**), indicating that acute loss of VEGFR2 signaling in ECs is not detrimental to adult islet vascular homeostasis or β cell proliferation. A similar observation was made previously when VEGF-A was acutely inactivated in adult β cells^18^. Next, we crossed VEGFR2^iΔEC^ mice with the existing βVEGF-A line to effectively perturb VEGF-A–VEGFR2 signaling in ECs at various time points during β cell loss and recovery. To control for any possible effects of Tm administration on compensatory β cell proliferation^27^, we treated all mice with Tm and designated Cre-negative (βVEGF-A; VEGFR2^fl/fl^) mice as controls to represent intact VEGFR2 signaling by ECs.

To determine the effect of proliferative ECs on MΦ recruitment and polarization, Tm was administered to knock down VEGFR2 (R2) in ECs prior to VEGF-A induction in β cells (**Figures 4A** and **S4A**). VEGF-A was induced in both control βVEGF-A; R2^fl/fl^ and βVEGF-A; R2^iΔEC^ genotypes, with efficient knockdown of VEGFR2 in the latter (**Figure S4B**). As expected, VEGF-A induction caused a 7% EC expansion and associated 16% β cell loss per islet area in βVEGF-A; R2^fl/fl^ controls, whereas EC and β cell area did not change in βVEGF-A; R2^iΔEC^ mice (**Figures 3B-D**). These results indicate that activation of VEGFR2 signaling in ECs by acute elevation of VEGF-A in the islet microenvironment is essential for islet hypervascularization and leads to β cell loss.

**Figure 3.**
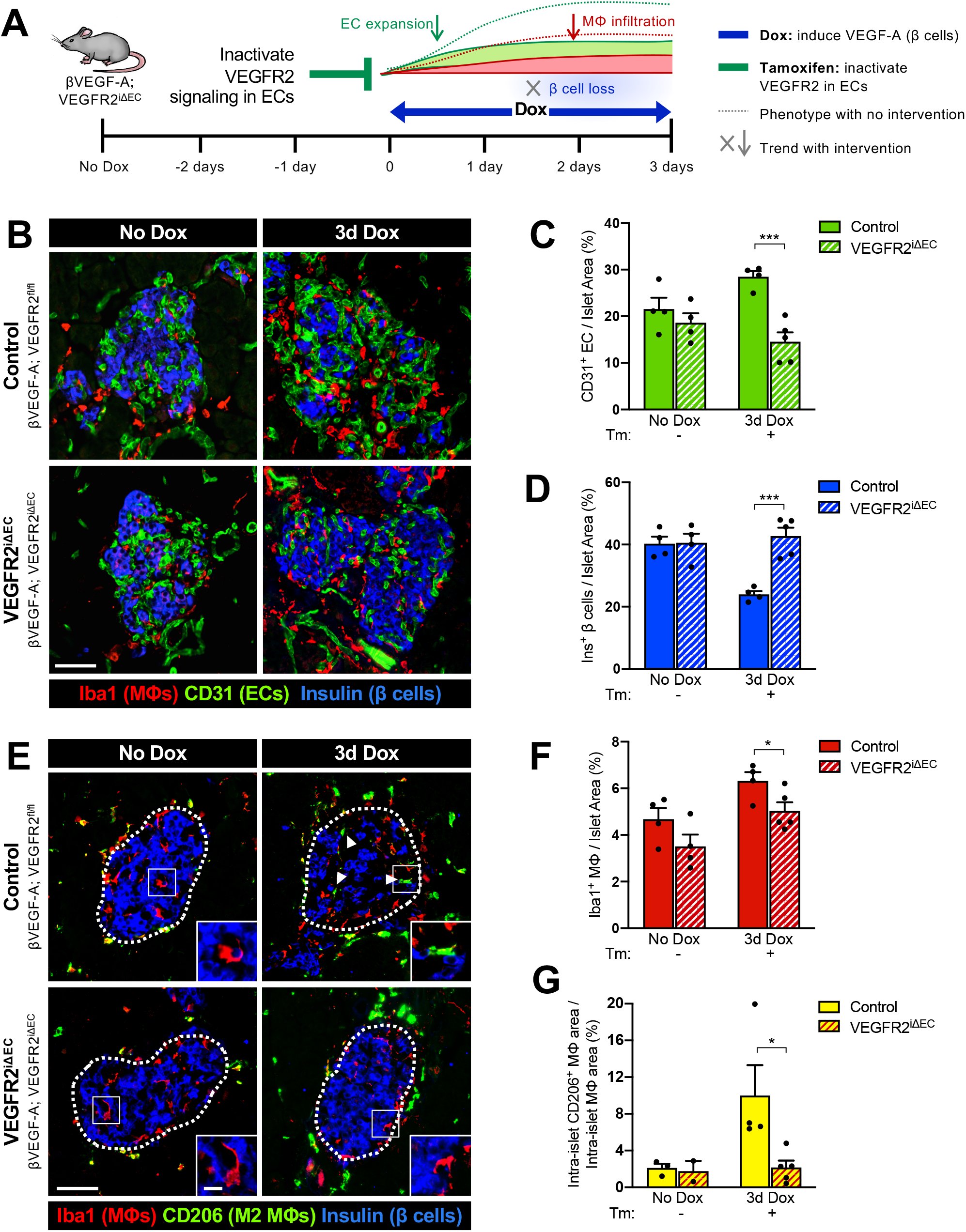
Inactivation of VEGFR2 signaling in endothelial cells prevents β cell loss and M2-like macrophage polarization by acute elevation of VEGF-A in the islet microenvironment. **(A)** To inactivate VEGFR2 in endothelial cells (ECs), control (βVEGF-A; VEGFR2^fl/fl^) and VEGFR2^iΔEC^ (βVEGF-A; VEGFR2^iΔEC^) mice were treated with Tamoxifen (Tm; 4mg s.c.) prior to VEGF-A induction. **(B)** Islet architecture displayed by labeling for macrophages (Iba1^+^), ECs (CD31^+^), and β cells (Ins^+^) at baseline (No Dox) and after 3d Dox. **(C–D)** Quantification (mean+SEM) of islet β cell and EC composition (10±1 × 10^5^ μm^2^ total islet area analyzed per animal). **(E)** Some intra-islet macrophages (MΦs) in control mice showed an “M2-like” phenotype (CD206^+^) after VEGF-A induction, indicated by arrowheads. Insets show representative intra-islet MΦs in each group and time point. **(F–G)** Quantification (mean±SEM) of MΦ infiltration and M2-like intra-islet MΦs (percent CD206^+^ Iba1^+^ of Iba1^+^), 3±2 × 10^5^ μm^2^ total islet area analyzed per animal. Each closed circle in bar graphs represents one animal. Asterisks indicate unpaired two-tailed t-tests between genotypes; *, p<0.05; ***, p<0.001. Scale bars in **(B)** and **(E)**, 50 μm; inset, 10 μm.

**Figure 4.**
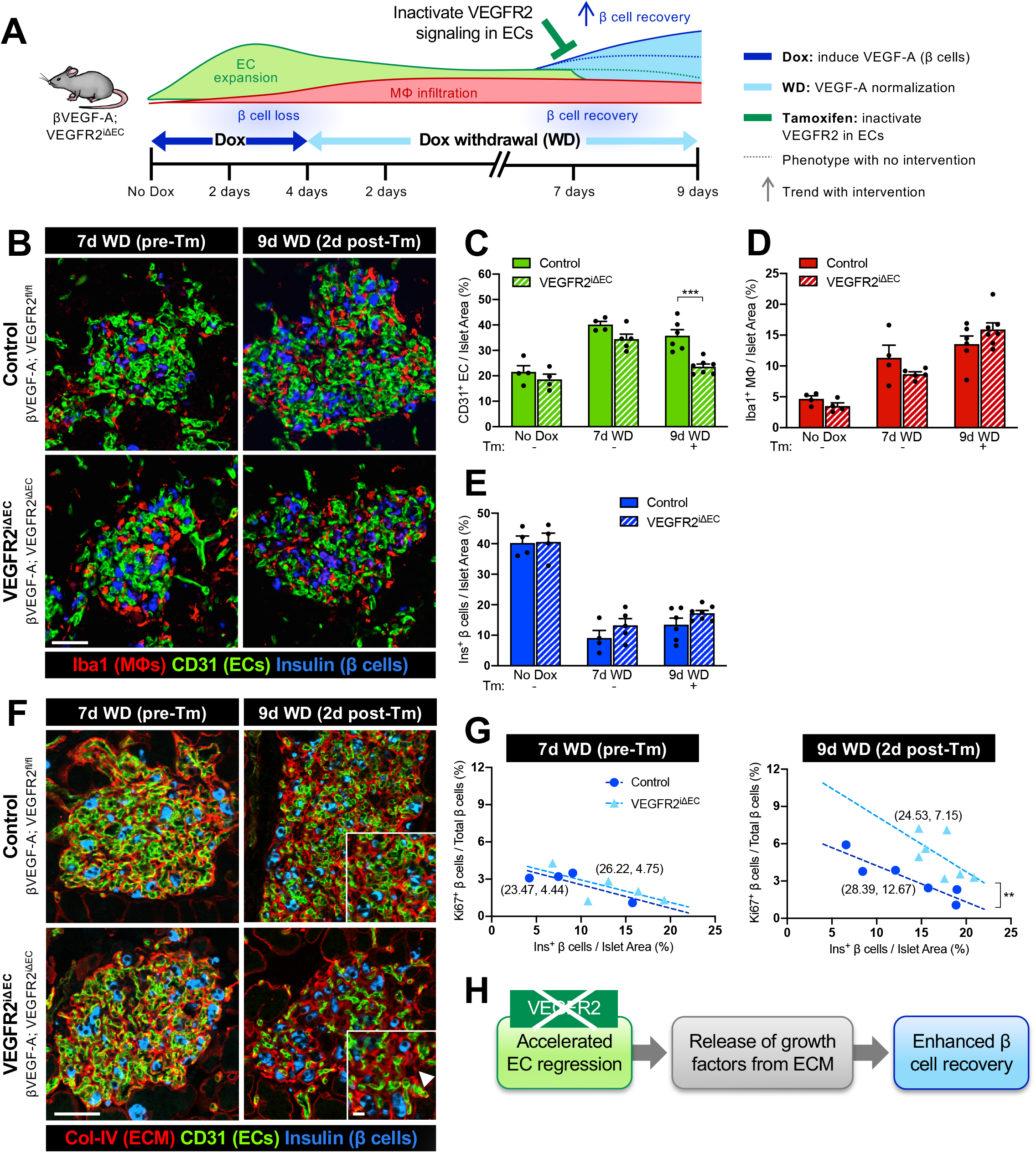
Phenotypic and structural changes to the islet microenvironment in response to VEGFR2 inactivation in quiescent endothelial cells during β cell recovery. **(A)** To inactivate VEGFR2 in endothelial cells (ECs) during β cell recovery, control (βVEGF-A; VEGFR2^fl/fl^) and VEGFR2^iΔEC^ (βVEGF-A; VEGFR2^iΔEC^) mice received Tamoxifen (Tm; 4mg s.c.) after 7 days (d) of Dox withdrawal (WD). **(B)** Islet architecture visualized by labeling for macrophages (Iba1^+^) ECs (CD31^+^, green), β cells (Insulin^+^, blue) at 7d WD (pre-Tm) and 9d WD (2d post-Tm). Scale bar, 50 μm. **(C–E)**Area quantification (mean±SEM) of islet ECs (**C**), MΦs (**D**), and β cells (**E**) by immunohistochemistry; 14±1 × 10^5^ μm^2^ total islet area analyzed per animal. Each circle represents one animal; asterisks indicate results of unpaired two-tailed t-tests between genotypes; ***, p<0.001. (**F**) Visualization of islet extracellular matrix by immunofluorescence (ECM; Col-IV^+^, red), ECs (CD31^+^, green), β cells (Insulin^+^, blue) at 7d WD (pre-Tm) and 9d WD (2d post-Tm). Scale bar, 50 μm; inset, 10 <m. Arrowhead in **(F)** points to ECM casts where ECs have regressed. **(G)** β proliferation rates (1,947±145 β cells per animal) plotted as a function of β cell loss (% β cells of total islet area) reveal a significant increase after VEGFR2 inactivation in quiescent ECs at 9d WD. Parentheses beside lines provide x- and y-intercepts derived from linear regression. At 9d WD, intercepts are significantly different; **, p<0.01. **(H)** Schematic of proposed mechanism for increased β cell area observed in VEGFR2^iΔEC^ mice.

Inactivation of VEGFR2 signaling in ECs significantly reduced, but did not completely prevent MΦ recruitment to βVEGF-A; R2^iΔEC^ islets compared to βVEGF-A; R2^fl/fl^ controls (5.0 vs. 6.3% per total islet area, respectively; p<0.05) (**Figures 3E** and **3F**). This suggests that MΦ infiltration can occur in response to VEGF-A alone, which is consistent with the fact that VEGF-A can mediate monocyte recruitment through VEGFR1 activity^28,29^. Still, the proliferative EC environment (intact VEGF-A–VEGFR2 signaling and β cell loss) leads to maximal MΦ recruitment. Interestingly, a subset of infiltrating MΦs in control islets expressed the M2-like marker CD206 (Mrc1), which is normally made only by exocrine MΦs^30,31^, while infiltrating MΦs in βVEGF-A; R2^iΔEC^ islets remained CD206^−^ (**Figures 3E** and **3G**). This observation suggests that VEGFR2 signaling in proliferative ECs and/or β cell loss promotes MΦ polarization to an M2-like phenotype.

### VEGFR2 inactivation in quiescent ECs accelerates EC regression, enhancing β cell recovery

To investigate the role of VEGFR2 signaling in quiescent ECs during β cell recovery, VEGF-A-mediated EC proliferation and β cell loss was first induced with 3-day Dox treatment in βVEGF-A; R2^iΔEC^ mice and controls, followed by 7 days of Dox withdrawal to allow VEGF-A to normalize and ECs to return back to quiescence (**Figures 4A, S5A**). Islet phenotype and β cell proliferation was assessed after 7 days of VEGF-A normalization (7d WD) and subsequently 2 days post-Tm treatment to inactivate VEGFR2 (9d WD). VEGF-A-induced EC expansion, β cell loss, and MΦ infiltration occurred in both genotypes, with no difference between the two groups at 7d WD prior to VEGFR2 inactivation (**Figures 4B-E**). As expected, VEGF-A expression continued to decline during Dox withdrawal and VEGFR2 was efficiently inactivated in βVEGF-A; R2^iΔEC^ mice within 48 hours of a single 4-mg Tm injection (**Figure S5B**). Surprisingly, unlike under normal homeostatic conditions (**Figure S3**), VEGFR2 inactivation in βVEGF-A; R2^iΔEC^ mice significantly accelerated islet EC regression compared to controls (**Figure 4D**) without changes in MΦ phenotype (**Figure S5C**) and number (**Figure 4C**), suggesting that neither MΦ retention nor polarization at this stage of β cell recovery are dependent on VEGFR2 signaling in quiescent ECs.

Since VEGFR2 inactivation in quiescent ECs of βVEGF-A; R2^iΔEC^ mice leads to a relatively rapid EC decline after Tm treatment (9d WD), we next evaluated the islet vascular regression by visualization of collagen IV, a major component of the islet ECM^32,33^ generated by intra-islet ECs. This study revealed that regressing islet capillaries leave behind vascular “casts” of ECM that are no longer associated with intact ECs (**Figure 4F,***9d WD*). Even more surprising was our finding that this accelerated decline in islet ECs enhances β cell proliferation at 9d WD (**Figure 4G**). We hypothesize that in contrast to β cell homeostasis, the regression of quiescent ECs during the β cell recovery phase stimulates release of growth factors from degraded ECM, thereby promoting β cell proliferation (**Figure 4H**).

### Identifying interactions between β cells, ECs, and MΦs in the βVEGF-A islet microenvironment

Our prior studies in the βVEGF-A model localized the stimulus for β cell proliferation to the islet microenvironment, ruling out any contribution from circulating factors that might reach the pancreas^1^. With key roles established for both MΦs and ECs in this system, we next sought to identify potential mechanisms and signaling pathways coordinating cell-cell and cell-matrix interactions. To do this we isolated βVEGF-A islets at baseline (No Dox) and during the course of VEGF-A induction (1wk Dox) and normalization (1wk WD) and purified islet populations of β cells, ECs, and MΦs at each of these time points for transcriptome analysis (**Figures 5A, S6A** and **S6B**).

**Figure 5.**
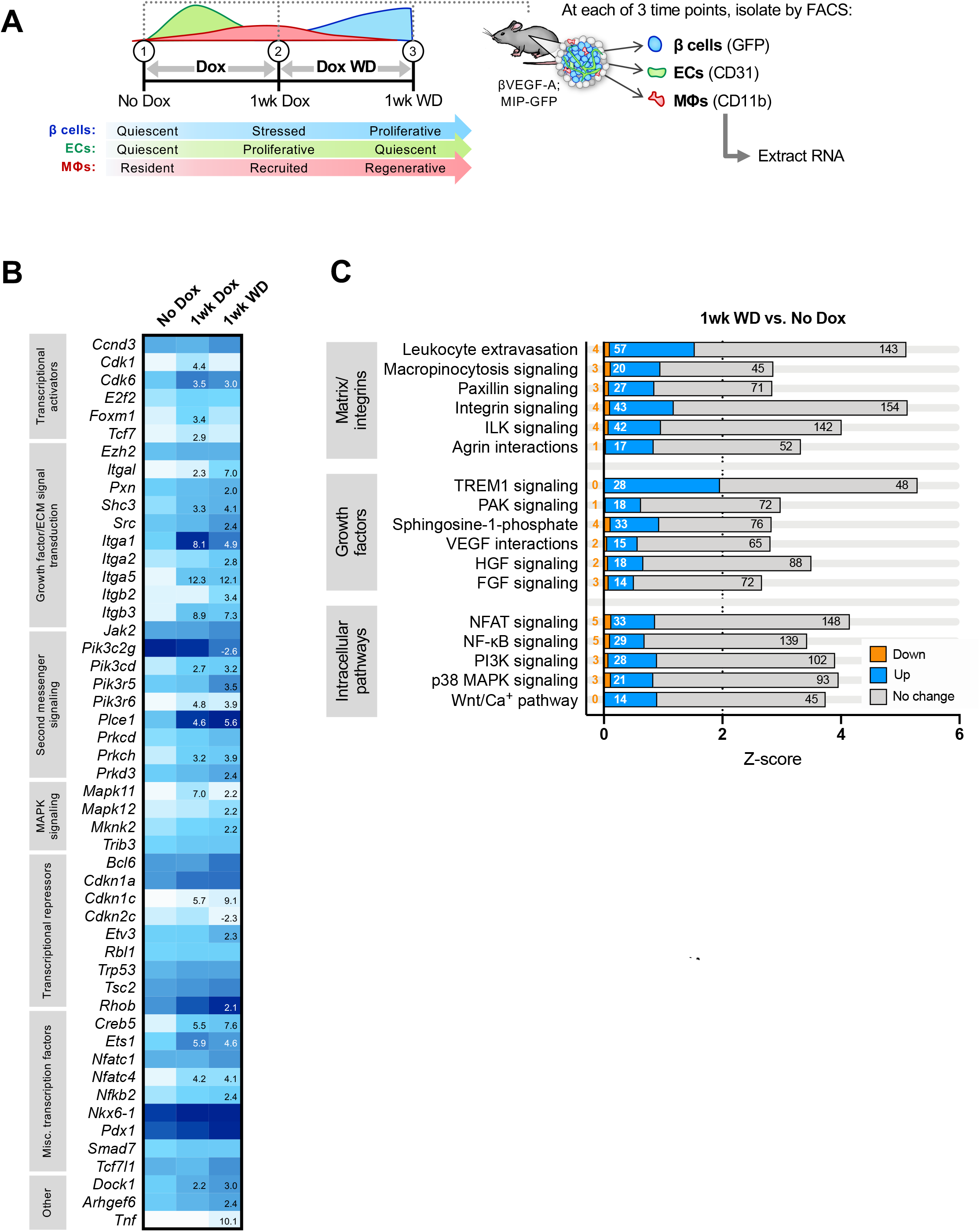
Genes and pathways upregulated in β cells during recovery reflect activation of intracellular signaling to promote proliferation. **(A)** Experimental schematic showing sorting of islet-derived β cells, endothelial cells (ECs), and macrophages (MΦs) from βVEGF-A mice at baseline (No Dox), during VEGF-A induction/β cell loss (1wk Dox), and during VEGF-A normalization/β cell recovery (1wk WD). See also **Figure S5B–E**. **(B)** Normalized expression of selected genes known to function in β cell proliferation^34–37^ at No Dox (n=4 replicates), 1wk Dox (n=5), and 1wk WD (n=3). Numbers listed in 1wk Dox and 1wk WD columns represent fold-change ≥2 or <−2 (p<0.05) as compared to No Dox. **(C)** Selected differentially regulated pathways during β cell recovery (1wk WD vs. No Dox), as determined by Ingenuity Pathway Analysis (IPA). Total bar width represents z-score; colored fractions indicate percentage of pathway genes that are up (blue), down (orange), and unchanged (grey). Number of genes per category is also listed within or adjacent to bars. A full list of significantly regulated pathways (z-score ≥2 or ≤−2, p<0.05) is provided in **Table S1**. FGF, fibroblast growth factor; HGF, hepatocyte growth factor; ILK, integrin-linked kinase; MAPK, mitogen-activated protein kinase; NFAT, nuclear factor of activated T cells; NF-kB, nuclear factor kappa-light-chain-enhancer of activated B cells; PI3K, phosphoinositide 3-kinase; PAK, p21-activated kinase; TREM1, triggering receptor expressed on myeloid cells 1; VEGF, vascular endothelial growth factor.

All of the nine sample types analyzed (3 cell types per each of 3 time points) demonstrate distinct transcriptional profiles, reflecting a high degree of uniqueness among three cell types and significant temporal changes in gene expression within each cell type (**Figures S6C-E**). By hierarchical clustering, the highest correlations were observed within one cell type across different time points; for example, β cells at No Dox are more similar to β cells at 1wk Dox and 1wk WD than they are to MΦs or ECs at any time point. Of the β cell samples, regenerative β cells (1wk WD) are more similar to stressed β cells (1wk Dox) than quiescent β cells (No Dox); in contrast, regenerative MΦs (1wk WD) are more similar to resident MΦs (No Dox) than to those recruited upon VEGF-A induction (1wk Dox), and quiescent ECs following VEGF-A normalization (1wk WD) are more similar to quiescent ECs at baseline (No Dox) than to proliferative ECs (1wk Dox) (**Figure S6D**).

With the induction of VEGF-A (1wk Dox), all cell populations increase expression of growth factors, matrix remodeling enzymes involved in tissue repair and matrix degradation (MMPs, ADAMs, ADAMTSs), as well as cell adhesion molecules involved in cell-matrix and cell-cell interactions (ICAM1, VCAM1, selectins), many of which remain elevated during β cell regeneration (1wk WD) (**Figure S7A**). Pathway analysis shows a high degree of cellular motility (**Figure S7B**), and extracellular organization and cell adhesion processes are significantly enriched in all cell types during β cell recovery (**Figure S8**).

Infiltrating MΦs show elevated expression of both pro-inflammatory and pro-regenerative markers during periods of initial monocyte recruitment (1wk Dox) and β cell regeneration (1wk WD). Pro-inflammatory genes like *Il12b* and *Il1a* are upregulated at 1wk Dox, while M2 markers such as *Ptgs1* and *Retnla* are upregulated and/or remained elevated at 1wk WD (**Figure S7A**). Upregulation of chemokines and cytokines by MΦs and corresponding increase in expression of receptors (e.g., *Cxcr4*, *Il12rb2*) by β cells suggests crosstalk between the two cell types and potential phenotypic effects on β cells in addition to MΦs. Similarly, increased growth factor expression in ECs and MΦs appears in concert with increased expression of growth factor receptors in β cells (**Figure S7A**).

At the peak of regeneration (1wk WD), β cells highly express integrins and other molecules that sense and respond to changes in the extracellular milieu (**Figure 5B**). Activation of integrin-mediated signaling and the integrin-linked kinase pathway were accompanied by increases in PI3K/Akt and MAPK signaling genes (**Figures S8A** and **5C**), many of which are known modulators of β cell proliferation^34–37^. Transcriptional activators *Cdk6* and *Foxm1* are initially upregulated more than 3-fold compared to baseline, followed by downregulation of suppressor *Cdkn2c* (p18) specifically during β cell recovery, consistent with previous studies of both mouse and human β cells^38–40^. *Creb5* and *Ets1* are notably increased as well. Collectively, these data provide evidence for a model where β cell proliferation is largely driven through MΦ–EC–β cell paracrine signaling, ECM remodeling, and cell-matrix interactions (**Figure 6**).

**Figure 6.**
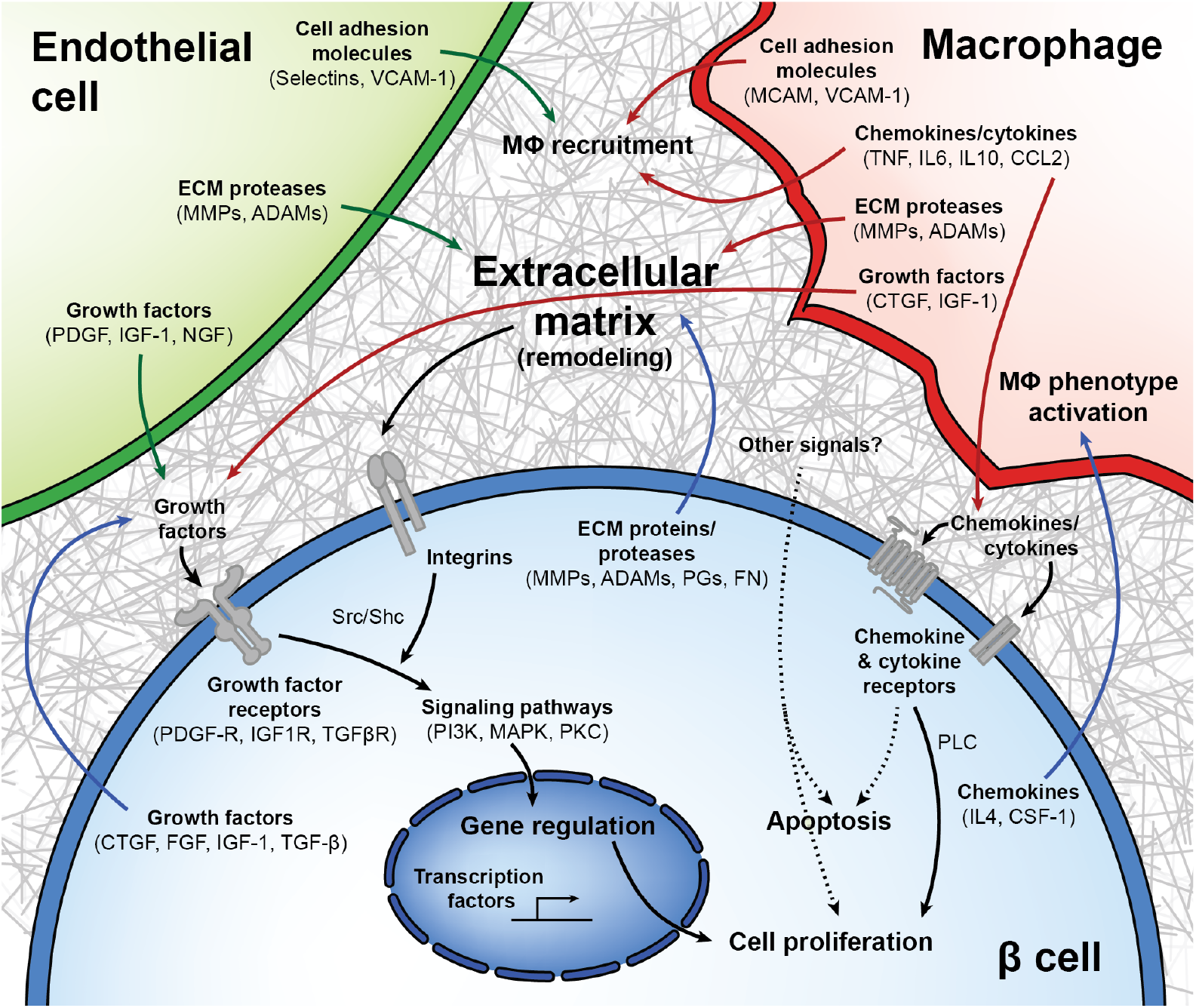
Model of interactions between β cells, macrophages, endothelial cells, and the extracellular matrix in β cell regeneration. Upon VEGF-A induction, intra-islet endothelial cells (ECs) proliferate while increasing expression of cell adhesion molecules and growth factors and altering their expression of integrins and extracellular matrix (ECM) remodeling enzymes. These adhesion molecules help recruit macrophages (MΦs), which upon islet infiltration also upregulate expression of cell adhesion molecules and pro- and anti-inflammatory chemokines and cytokines, influencing further MΦ recruitment in addition to signaling through chemokine and cytokine receptors on β cells. Chemokine and cytokines become increasingly less inflammatory as VEGF-A normalizes, as MΦs produce growth factors and matrix remodeling enzymes that may promote β cell proliferation. Upon VEGF-A induction, β cells exhibit enrichment for several integrin pathways and other proteins involved in ECM remodeling and cell-matrix interactions in addition to regulating expression of chemokines known to support a regenerative (M2 or alternative) MΦ phenotype. Growth factors from all cell types act on an increased number of growth factor receptors being expressed on β cells, activating downstream signals converging on the PI3K/Akt and MAPK pathways. Other signals from cells in the microenvironment, or from the rapidly remodeling ECM, may also play a role in β cell proliferation.

## DISCUSSION

Adult β cells have a very limited proliferative capacity and signals regulating this process are incompletely understood. The complexity of the islet microenvironment and difficulty modeling this complexity *in vitro* often constrains investigation of β cell proliferation to reductionist approaches, removing components of the *in vivo* environment known to contribute to the establishment and maintenance of β cell mass and thereby limiting the utility of these studies. Here we utilized the βVEGF-A mouse model^1^ in which signals from the local islet microenvironment, including β cells, ECs, and MΦs, promote proliferation of human as well as mouse β cells. To dissect how these microenvironmental components influence β cell loss and recovery within the context of this complex and dynamic *in vivo* model, we developed new experimental tools to intricately modulate EC and MΦ populations and their signaling within the islet. These tools, coupled with transcriptional analysis of temporal changes in gene expression patterns of β cells, ECs, and MΦs, allowed us to gain important insight into the cell-cell and ECM-mediated signaling in this model, which work in concert to promote β cell proliferation (**Figure 6**).

### Role of macrophages

Our transcriptome analysis suggested that MΦ recruitment to islets is facilitated not only by VEGF-A signaling but also by increased expression of cell adhesion molecules on both MΦs and ECs, as well as increased production of chemokines and cytokines from MΦs already in the islet microenvironment^41^. By specifically depleting MΦs in βVEGF-A mice using clodronate liposomes, we demonstrated that MΦs are required for β cell regeneration in the islet microenvironment. These recruited MΦs have a unique hybrid phenotype: they express markers of both classical pro-inflammatory (M1) activation as well as alternative (M2) activation promoting tissue repair and regeneration. Though MΦ-derived cytokines and chemokines can exacerbate β cell stress^42,43^, the comparable β cell loss in MΦ-depleted and control islets suggests that MΦs do not promote β cell apoptosis in this model. Instead, MΦs downregulate key pro-inflammatory cytokines (*e.g.*, *Tnf, Il6*) at 1wk WD, concomitant with β cell proliferation. Their phenotypic profile at this stage is most reminiscent of phenotyping subtypes “M2a” (high expression of scavenger and phagocytic receptors like Retnlb/Fizz2, Ym1, and Mrc1; secretion of profibrotic and trophic factors like fibronectin, IGF, and TGFβ) and “M2c” (associated with removal of apoptotic cells)^44–46^. This phagocytic phenotype is consistent with our finding that MΦs at this time point (1wk WD) likely have an effect on islet remodeling and composition separate from their effect on β cell proliferation. Together, these data suggest a phenotypic shift in MΦs that facilitates β cell recovery and return to homeostasis in the islet, in line with an increasing number of studies observing tissue-restorative effects of MΦs triggered in various β cell injury models^11,12,47^. Further studies will be necessary to clarify the specific signals that regulate the phenotypic shift and define the cues that govern their withdrawal.

### Role of endothelial cells

Regulated EC-derived signals are necessary for pancreatic and islet development^17,48–50^, optimal adult β cell function^18^, and islet revascularization after transplantation^51–54^. In addition, ECs and VEGFR2 signaling have been previously implicated in organ regeneration^13,16^. For example, during liver regeneration, sinusoidal ECs appear to have a biphasic effect in mediating hepatic reconstitution; proliferative angiogenesis of sinusoidal ECs inhibits hepatocyte self-renewal, whereas quiescent sinusoidal ECs stimulate hepatic regeneration^13^. To determine the role of proliferative and quiescent ECs in β cell loss and recovery, we generated a compound βVEGF-A model with inducible VEGFR2 inactivation (βVEGF-A; VEGFR2^iΔEC^) that allowed us to modulate VEGF-A–VEGFR2 signaling in ECs.

Our studies show that intact VEGFR2 signaling is required for the proliferative angiogenesis of islet endothelium, which ultimately results in β cell loss. Although transcriptome analysis did not reveal specific signals produced by proliferating ECs that would lead to β cell apoptosis during VEGF-A induction, there are significant changes in expression of matrix remodeling enzymes and cell adhesion molecules. Based on the importance of cell-matrix interactions in β cell survival *in vitro*^20,55^, we hypothesize that ECM changes associated with rapidly expanding endothelium likely contribute to β cell loss in the βVEGF-A system. In addition, we show that intact VEGFR2 signaling is necessary for accumulation of intra-islet MΦs expressing CD206, a marker of an M2 phenotype. This observation is consistent with MΦ activation by EC-derived cues in the context of acute injury, a phenomenon that has been noted in numerous tissues^56–58^. Furthermore, a similar M2 phenotype (high IL-10, CD206) was shown to be required for β cell regeneration after DT-mediated ablation^12^, and elsewhere CD206 has been linked to increased expression of *Tgfb1* and *Egf* ^59^ and promotion of β cell proliferation via a Smad7-cyclin pathway^11^.

In contrast to lung and liver regeneration^13,16^, VEGFR2 inactivation in quiescent ECs (1wk WD) resulted in accelerated β cell recovery, which is associated with rapid islet capillary regression, leaving behind vascular “casts” of ECM components. We postulate that β cell proliferation is promoted by release of growth factors from degraded ECM. Though ECs are no longer expanding at the 1wk WD time point, islets still contain quite extensive capillary networks that might be sustained by the residual extracellular VEGF-A we observed, as well as possibly other growth factors. Interestingly, VEGFR2 inactivation during β cell homeostasis does not impact β cell proliferation and islet capillary morphology. It will be important to assess the role of quiescent ECs and VEGFR2 signaling in other β cell injury models in the future. In addition, these findings highlight that ECs have different roles in different tissues and during various physiologic and pathologic stressors.

### Role of extracellular matrix and integrated cellular interactions in β cell proliferation

By deconstructing the contributions of the cellular components of the islet microenvironment of βVEGF-A mice, we were able to uncover a role for ECM remodeling and signaling in β cell proliferation. MΦs, ECs, and β cells all show transcriptional regulation of integrin receptors and ECM remodeling enzymes, some of which have been shown previously to affect β cell proliferation^22,60–62^. In addition to shaping cell-matrix signaling, ECM reorganization can also lead to the release and/or activation of matrix-sequestered growth factors^63–66^. This is particularly relevant given the temporal increase of growth factor receptor expression in β cells during VEGF-A induction and normalization, suggesting heightened β cell responsiveness to signals from the rapidly remodeling ECM as VEGF-A normalizes. Together, these data provide evidence that ECM-bound growth factors released during EC regression promote β cell proliferation.

Remodeling of extracellular milieu can also influence MΦ phenotype^67,68^, though more work is required to define specific signaling pathways that regulate this process in the pancreas. However, our transcriptome data provides evidence that both MΦs and ECs are quite attuned to their rapidly changing environment, with integrins and cell adhesion molecules being some of the most dynamically regulated genes in both cell types. Changes in these molecules are likely regulating MΦ recruitment and/or polarization^69–72^, which may explain the phenotypic shift that happens between MΦ recruitment (occurring rapidly in response to VEGF-A) and the appearance of β cell proliferation (during VEGF-A normalization).

It is also possible that β cells undergo intrinsic changes heightening their sensitivity to extracellular signals, supported by the observation that ECs and MΦs increase expression of several growth factors known to promote β cell proliferation (IGF-1, PDGF, and CTGF)^37,73–75^ while β cells simultaneously upregulate expression of corresponding receptors (*Igf1r*, *Pdgfr*).

Signaling cascades activated by integrins and growth factors exhibit extensive downstream crosstalk and protein activity that makes it difficult to determine pathway activation status based solely on gene expression. Nonetheless, we did observe transcriptional changes to components of the PI3K/Akt, PLC, and MAPK pathways, as well as upregulation of transcription factors regulated by the MAPK pathway^34–37^ suggesting that the cell-cell and cell-ECM interactions may converge on the activation of these pathways leading to β cell proliferation. We therefore propose a model in which coordination of growth factors – whose bioavailability is likely modulated by ECs and MΦs – together with increased integrin signaling promotes activation of pro-proliferative pathways in surviving β cells during VEGF-A normalization to ultimately restore β cell mass (**Figure 6**).

Recognizing the key role of cellular and extracellular components of the islet microenvironment on β cell development, function, and homeostasis is critical to further our understanding of signals regulating adult β cell proliferation. Islet microenvironmental signaling in the βVEGF-A system promotes human β cell proliferation^1^, which prompted us to develop new strategies to disentangle the roles of various microenvironmental components in this regenerative process. However, moving forward it will be important to further explore the mechanisms and combination of specific cytokines, growth factors, and other microenvironmental signals that activate and regulate relevant mitogenic signaling pathways in human β cells. Overall, these studies highlight the importance of developing innovative approaches to examine β cells *in vivo* in order to more completely understand the complex microenvironmental factors regulating β cell function and regeneration.

## METHODS

### Mouse Models

All animal studies were approved by the Institutional Animal Care and Use Committee at Vanderbilt University Medical Center, and animals were kept in facilities monitored by the Vanderbilt University Division of Animal Care on a 12 hour light/12 hour dark schedule with unrestricted access to standard chow and water. Mouse models and abbreviations used to describe them are summarized in **Table S2**.

#### RIP-rtTA; TetO-VEGF (βVEGF-A) mice

The original bitransgenic mice with doxycycline (Dox)-inducible β-cell-specific overexpression of human VEGF-A_165_ (abbreviated βVEGF-A) were generated by crossing RIP-rtTA male mice and TetO-VEGF female mice, both on a C57BL/6 background^76–80^. These mice were generously provided by Dr. Shimon Efrat of Tel Aviv University and Dr. Peter Campochiaro of Johns Hopkins University, respectively. In this βVEGF-A model the rat *Ins2* promoter drives expression of the tetracycline-responsive rtTA transactivator specifically in pancreatic β cells. Upon exposure to Dox, the rtTA transactivator binds the tetracycline operator (*TetO*), driving expression of human VEGF-A_165_ in β-cells. Details of Dox preparation and administration are included below.

#### Cd5-CreER; VEGFR2^fl/fl^ (VEGFR2^iΔEC^) mice

Mice with Tamoxifen (Tm)-inducible EC-specific knockout of VEGFR2 (abbreviated VEGFR2^iΔEC^) were generated by crossing Cd5-CreER male mice and VEGFR2^fl/fl^ female mice (see **Table S3**, crosses A1-A2). Heterozygous VEGFR2^fl/wt^ mice on a C57BL/6 background were obtained from Jackson Laboratories (stock #018977) and bred to create a homozygous VEGFR2^fl/fl^ line. Frozen sperm from the Cd5-CreER line^26^ was generously provided by Dr. Yoshiaki Kubota of Keio University, and *in vitro* fertilization (IVF) was performed by the Vanderbilt Genome Editing Resource using female C57BL/6 mice (Jackson Laboratories, stock #000664). In the VEGFR2^iΔEC^ model *loxP* sites were inserted flanking VEGFR2 exon 3, and the *Cdh5* (*VE-cadherin*) promoter drives expression of Tm-inducible Cre recombinase in vascular ECs. Upon exposure to Tm, Cre recombinase translocates to the nucleus, excising VEGFR2 exon 3 through Cre-*loxP* recombination and subsequently preventing VEGFR2 expression in ECs. Details of Tm preparation and administration are described below.

#### βVEGF-A; VEGFR2^iΔEC^ mice

To generate an inducible model of EC-specific knockdown of VEGFR2 in βVEGF-A mice, Cd5-CreER and VEGFR2^fl/fl^ mice were crossed with RIP-rtTA and TetO-VEGF transgenic mice as outlined in **Table S3**. The final cross produced both βVEGF-A; VEGFR2^iΔEC^ mice as well as Cre-negative sibling controls (βVEGF-A; VEGFR2^fl/fl^). Due to the inefficient induction of VEGF-A in female mice (the single copy of the *TetO* transgene is subject to × chromosome inactivation), only male mice were used for experiments.

#### βVEGF-A; MIP-GFP mice

To enable fluorescence-activated cell sorting (FACS) of pancreatic β cells from βVEGF-A mice, a transgene was separately introduced into the βVEGF-A mouse model by crossing MIP-GFP mice on a C57BL/6 background^81^ (Jackson Laboratories, stock #006864) with RIP-rtTA and Tet-O-VEGF-A mice (abbreviated βVEGF-A; MIP-GFP). In these mice, the mouse *Ins1* promoter drives green fluorescent protein (GFP) expression in β cells.

#### DNA extraction and genotyping

Mouse models used in our breeding schemes were maintained by genotyping using the primers and PCR conditions listed in **Table S4**. DNA was extracted and PCR reactions were performed with tail snips from mice as described previously^1^. Thermal cycler conditions listed in **Table S4** were used to amplify DNA before resolving on agarose gels with 100 ng/ml ethidium bromide in 1X Tris/Borate/EDTA (TBE) buffer as indicated.

#### Compound preparation and administration

VEGF-A transgene expression was activated in βVEGF-A mice by Dox administration (5 mg/ml) in light-protected drinking water containing 1% Splenda^®^ for a period of 3-7 days. VEGFR2 knockdown was induced in VEGFR2^iΔEC^ mice by subcutaneous injection of 4 mg Tm (20 mg/ml; 200 μl). Tm (20 mg/ml) was prepared fresh in filter-sterilized corn oil the day before each injection and allowed to dissolve overnight on a shaker at room temperature, protected from light. Vetbond tissue adhesive (3M) was used to seal injection sites to prevent oil leakage. Clodronate-mediated macrophage depletion in βVEGF-A mice was accomplished by injecting 150-200 μl clodronate liposomes (5 mg/ml; Clodrosome) retro-orbitally every other day for a 1-2 week period (4-8 total injections). Liposome injections began one day before Dox administration and continued for 1 week after in mice harvested at later time points. Control liposomes with the same lipid composition (Clodrosome) were administered to βVEGF-A mice using the same route, volume, and schedule. Mice receiving liposome injections were supplemented with Transgenic Dough Diet (21.2% protein, 12.4% fat, 46.5% carbohydrate; BioServ) throughout the course of the experiment.

#### Glucose measurements

Random (non-fasted) plasma glucose levels were measuring by obtaining whole blood from nicked tail veins using an Accu-chek glucose meter (Roche Diagnostics) calibrated according to the manufacturer’s instructions.

### Tissue Collection and Fixation

Mouse pancreata were collected from anesthetized mice prior to cervical dislocation. Organs were washed in ice-cold 10 mM phosphate buffered saline (PBS), then fat and other excess tissue was removed before pancreata were weighed and processed. Fixation was performed in 0.1 M PBS containing 4% paraformaldehyde (Electron Microscopy Sciences) for 2-3 hours on ice with mild agitation, then organs were washed in four changes of 0.1 M PBS over 2 hours and equilibrated in 30% sucrose/0.01 M PBS overnight. After blotting to remove excess sucrose, tissues were mounted in Tissue Tek cryomolds filled with Tissue-Plus Optimal Cutting Temperature (OCT) compound (VWR Scientific Products). Tissue molds were placed on dry ice until the OCT was set, then stored at −80°C. Tissues were sectioned from 5-10 μm thick on a Leica CM1950 cryostat (Leica) and these cryosections were attached to Superfrost Plus Gold slides (ThermoFischer Scientific).

### Immunohistochemistry, Imaging, and Analysis

Immunohistochemical analysis was performed on serial 8-10 μm pancreatic cryosections as described previously^1,18^. Briefly, tissue permeabilization was conducted using 0.2% Triton-X in 10nM PBS and blocking using 5% normal donkey serum in 10mM. Primary and secondary antibody incubations (listed in **Table S5**) were performed in buffer with 0.1% Triton-X and 1% BSA and nuclei were counterstained with DAPI. Slides were mounted using SlowFade Gold antifade reagent (Invitrogen Molecular Probes) and sealed with fingernail polish prior to imaging.

Digital images were acquired with a Leica DMI6000B fluorescence microscope equipped with a Leica DFC360FX digital camera (Leica), a laser scanning confocal microscope (Zeiss LSM510 META or LSM880, Carl Zeiss), and a ScanScope FL (Aperio). Image analysis was performed using MetaMorph 7.7 software (Molecular Devices), ImageScope software (Aperio), or HALO software (Indica Labs).

For analysis of islet composition, images of entire pancreatic sections were captured at 20x magnification using a ScanScope FL system. Islet area was annotated manually based on insulin staining, and HALO algorithms were used to calculate area of β cells (Insulin^+^), ECs (CD31^+^), and MΦs (Iba1^+^). For β cell proliferation, cells were deemed positive for Ki67 only when at least 75% of the nucleus was surrounded by insulin^+^ cytoplasm.

### Flow Cytometry and Cell Sorting

Flow analysis and sorting was performed in collaboration with the Vanderbilt Flow Cytometry Core. Peripheral blood (50-100 μl) was collected from the retro-orbital sinus of βVEGF-A mice using heparinized capillary tubes 24 hours after beginning liposome injections to evaluate depletion of circulating monocytes. Blood was incubated for 3-5 minutes at 37°C with 1 ml warmed, filter-sterilized erythrocyte lysis buffer (8.26 g ammonium chloride, 1 g potassium bicarbonate, and 0.38 g EDTA in 1 L Milli-Q water). Cells were pelleted by centrifuging at 1800 rpm for 2-3 minutes at 4°C and supernatant discarded. Incubation with erythrocyte lysis buffer was repeated, and then cells were washed with 1 ml FACS buffer (2 mM EDTA and 2% FBS in 10 mM PBS) prior to antibody incubation. Blood from WT mice was collected for antibody compensation controls.

Isolated islets from βVEGF-A; MIP-GFP mice handpicked in Clonetics EGM MV Microvascular Endothelial Cell Growth Medium (Lonza) were washed 3 times with 2 mM EDTA in 10 mM PBS and then dispersed by incubating with Accutase (Innovative Cell Technologies) at 37°C for 10 minutes with constant pipetting. Accutase was quenched with EGM MV media, and then islet cells were washed twice with the same media and counted using a hemocytometer prior to antibody incubation. Anti-rat Ig, κ CompBead Plus Compensation Particles (BD Biosciences) and EasyComp Fluorescent Particles, GFP (Spherotech) were used as single color compensation controls for islet cell sorts.

Peripheral blood and islet cells prepared as described above, and anti-rat Ig compensation particles were incubated for 15-20 minutes at 4°C with fluorophore-conjugated antibodies in FACS buffer followed by one wash with FACS buffer. All antibodies for flow cytometry and their working dilutions are listed in **Table S5**. Prior to analysis or sorting, either propidium iodide (0.05 μg/100,000 cells; Invitrogen Molecular Probes) or DAPI (0.25 μg/1,000,000 cells; Invitrogen Molecular Probes) was added to samples for non-viable cell exclusion. Flow analysis was performed using an LSRFortessa cell analyzer (BD Biosciences) and a FACSAria III cell sorter (BD Biosciences) was used for FACS. Analysis of flow cytometry data was completed using FlowJo 7.6.5-10.2.1 (FlowJo LLC).

### RNA Isolation, Sequencing, and Analysis

Sorted islet-derived cells (8,000-400,000/sample) were added to 200-400 μl lysis/binding solution in the RNAqueous micro-scale phenol-free total RNA isolation kit (Ambion). Trace contaminating DNA was removed with TURBO DNA-free (Ambion). RNA quality control quantification was performed using a Qubit Fluorometer (Invitrogen, Carlsbad, CA) and an Agilent 2100 Bioanalyzer. All RNA samples had an RNA integrity number (RIN) ≥5.0. RNA was amplified using the Ovation system (NuGen Technologies) according to standard protocol. Amplified cDNA was sheared to target 300bp fragment size and libraries were prepared using NEBNext DNA Library Prep (New England BioLabs). 50bp Paired End (PE) sequencing was performed on an Illumina HiSeq 2500 using traditional methods^**82,83**^. Raw reads were mapped to the reference mouse genome mm9 using TopHat v2.0^**84**^and aligned reads were then imported onto the Avadis NGS analysis platform (Strand Scientific). Transcript abundance was quantified using the TMM (Trimmed Mean of M-values) algorithm^**85,86**^. Samples were compared by principle component analysis (PCA) and hierarchal clustering analysis. A minimum expression cutoff (normalized expression ≥20 at one or more time points) was applied before determining differential expression between samples, which was calculated on the basis of fold change (cutoff ≥2 or ≤−2) with p-values estimated by z-score calculations (cutoff 0.05) as determined by the Benjamini Hochberg false discovery rate (FDR) method^**87**^. Differentially expressed genes were further analyzed through Ingenuity Pathway Analysis (IPA, Qiagen) and Gene Ontology (GO) analysis using DAVID^**88**^. RNA quality control, amplification, sequencing, and analysis were performed in collaboration with the Genomic Services Laboratory at HudsonAlpha Institute for Biotechnology.

### Statistical Analysis

Prism software (GraphPad) was used to perform all statistical analyses for immunohistochemistry. In all experiments manipulating MΦs and ECs, control and experimental groups were compared at each time point using an unpaired t test. For analysis of proliferative ECs, unpaired t tests were also used to compare baseline (No Dox) and VEGF-A induction (3d Dox) time points within each group. For analysis of quiescent ECs, a one-way analysis of variance (ANOVA) was used to analyze time points within each group, followed by Tukey’s multiple comparison test to compare each time point with baseline (No Dox). Unless otherwise noted, data are expressed as mean + standard error of mean (SEM). Statistical analysis of RNA-sequencing data is described above (see RNA Isolation, Sequencing, and Analysis).

## AUTHOR CONTRIBUTIONS

Conceptualization, D.C.S., K.I.A., M.B., and A.C.P.; Methodology, D.C.S., K.I.A., M.B., and A.C.P.; Investigation, D.C.S., K.I.A., T.M.R., A.H., Z.K., R.A., G.P., R.J., and M.B.; Formal Analysis, N.P.; Writing – Original Draft, D.C.S., M.B., and A.C.P.; Writing – Review & Editing, all authors; Funding Acquisition, A.C.P.; Supervision, A.C.P., M.B., and S.E.L.

## ACKNOWLEDGEMENTS

We thank Drs. Y. Kubota, S. Efrat, P. Campochiaro, and M. Gannon for providing mice. We are grateful to Drs. A. Pozzi, A. Hatzopoulos, D. Jacobson, R. Stein, P. Kendall, J. Thomas, V. Babaev, P. Young, and Y. Dor for helpful discussions. This work was supported by grants from the Department of Veterans Affairs, the JDRF, the NIH (DK106755, DK89572, DK66636, DK69603, DK63439, DK62641, DK72473, DK94199, DK68764, DK97829, DK11232, DK117147, DK104211), and the Vanderbilt Diabetes Research and Training Center (DK20593). Islet isolation was performed in the Vanderbilt Islet Procurement and Analysis Core (DK20593). Image acquisition was performed in part through use of the Vanderbilt Cell Imaging Shared Resource (CA68485, DK20593, DK58404, DK59637, EY08126) and Vanderbilt Islet Procurement and Analysis Core (DK20593). Flow cytometry was performed in the Vanderbilt Flow Cytometry Shared Resource (P30 CA68485, DK058404). Cd5-CreER mouse line rederivation was performed by the Vanderbilt Genome Editing Resource of the Center for Stem Cell Biology (CA68485, DK20593).

## DATA & RESOURCE AVAILABILITY

Authors are in the process of submitting RNA sequencing data to the Gene Expression Omnibus (GEO) database of the National Center for Biotechnology Information (NCBI). Additional datasets and materials generated during the current study are available from the corresponding author on reasonable request.

## SUPPLEMENTAL FIGURES

**Figure S1.**
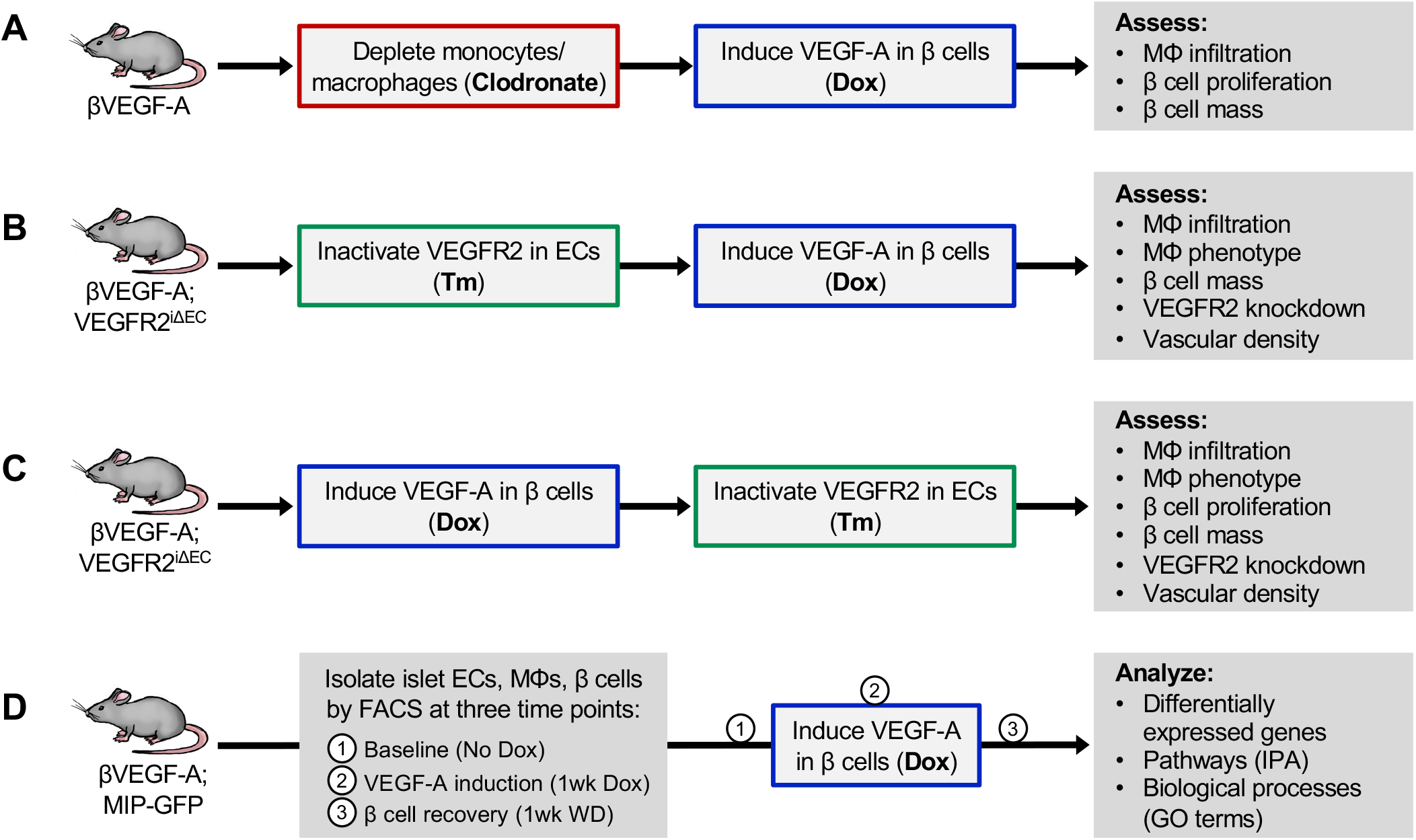
Schematic representation of experiments to determine the role of macrophages and endothelial cells in the βVEGF-A mouse model. **(A)** Clodronate liposomes were used to suppress macrophage (MΦ) infiltration, followed by administration of doxycycline (Dox) to overexpress VEGF-A in β cells. **(B–C)** To modulate endothelial cells (ECs), an inducible EC-specific VEGFR2 knockout mouse model Cd5-CreER; VEGFR2^fl/fl^ (VEGFR2^iΔEC^)^26^ was bred into the βVEGF-A line. Tamoxifen (Tm) was used to inactivate VEGFR2 in ECs either prior to **(B)** or after **(C)** VEGF-A induction to discern the role of proliferative and quiescent ECs, respectively in the β cell loss and recovery. **(D)** ECs, MΦs, and β cells were isolated from βVEGF-A mice at three time points; introduction of the MIP-GFP transgene facilitated β cell sorting. For additional details, see **Methods**.

**Figure S2.**
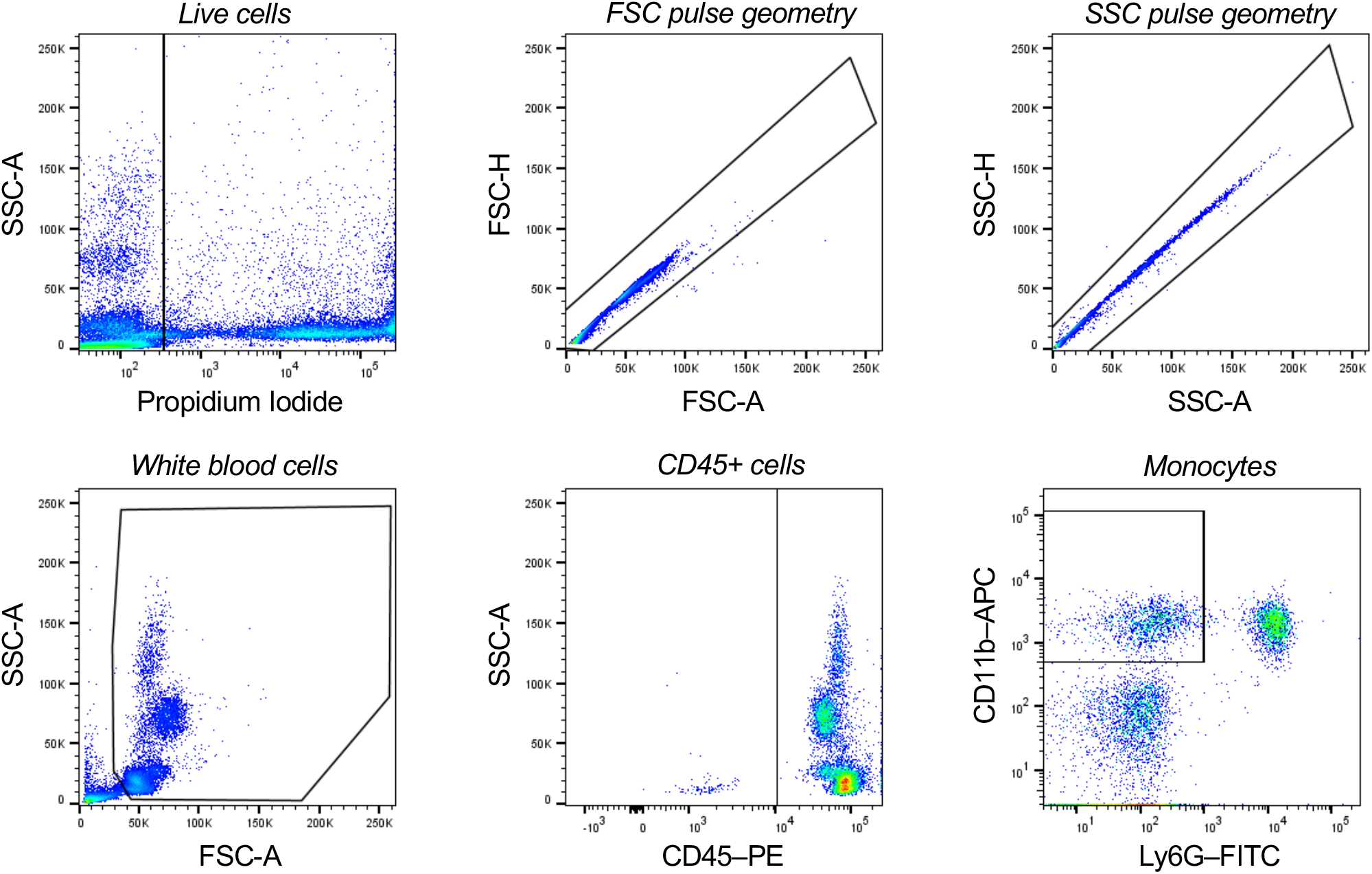
Gating strategy for flow cytometry analysis of circulating monocytes. Peripheral blood (50-100 μl) was collected from the retro-orbital sinus of βVEGF-A mice to assess monocyte depletion following clodronate treatment. Cell debris were excluded by forward scatter (FSC) and side scatter (SSC), single cells were identified by voltage pulse geometry, and non-viable cells were excluded using propidium iodide. From remaining white blood cells, CD45 was used as a pan-leukocyte marker and monocytes were identified by a CD11b^+^ Ly6G^−^ profile. Approximately 10,000 white blood cells were analyzed and quantified per each animal. See also **Figure 2B**.

**Figure S3.**
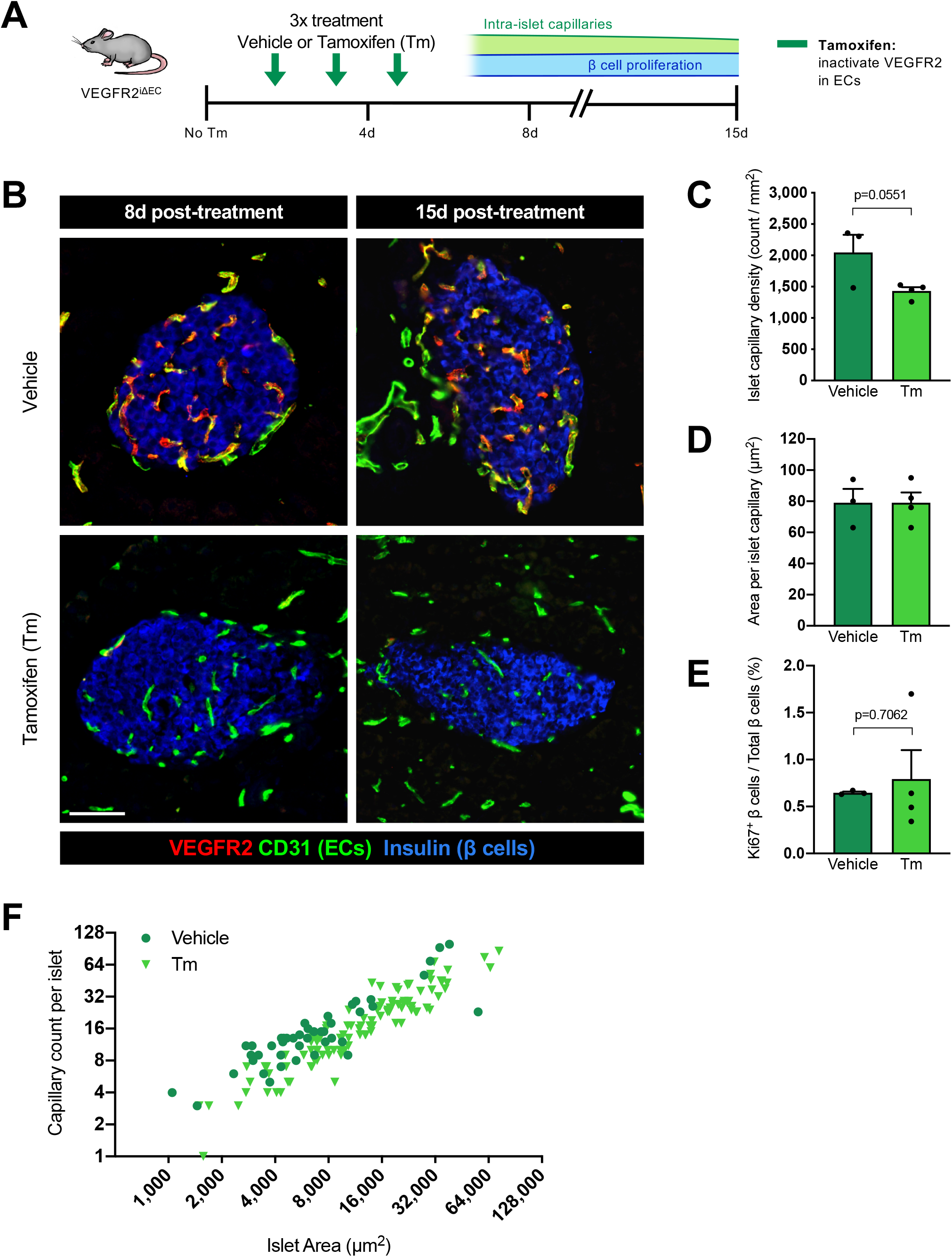
Acute ablation of VEGFR2 in endothelial cells does not alter islet vascular homeostasis or β cell proliferation. **(A)** To inactivate VEGFR2 in endothelial cells (ECs), Cdh5-CreER^T2^; VEGFR2^fl/fl^ (VEGFR2^iΔEC^) mice received treatments (Tx) of corn oil (vehicle) or tamoxifen (Tm; 4mg s.c.) every other day for 5 days. **(B)** Representative images of islet ECs (CD31^+^) and β cells (Insulin^+^) in vehicle- and Tm-treated VEGFR2^iΔEC^ mice. VEGFR2 expression (red) is virtually undetectable in ECs at 8 or 15 days (d) after initial Tx. Scale bar, 50 μm. **(C–E)** Quantification (mean±SEM) of islet architecture in VEGFR2^iΔEC^ mice at 8d post-Tx; each circle represents one animal. Islet capillary density **(C)** and area per islet capillary **(D)** determined by CD31^+^ and Insulin^+^ stain; 138±15 × 10^5^ μm^2^ total islet area analyzed per animal. **(E)** Basal β cell proliferation rate; 1,892±259 cells counted per animal. Statistics shown in **(C)** and **(E)** reflect unpaired two-tailed t-tests. **(F)** In both vehicle- and Tm-treated mice, the number of islet capillaries increases proportionately with islet size. Note both axes have a base 2 logarithmic scale.

**Figure S4.**
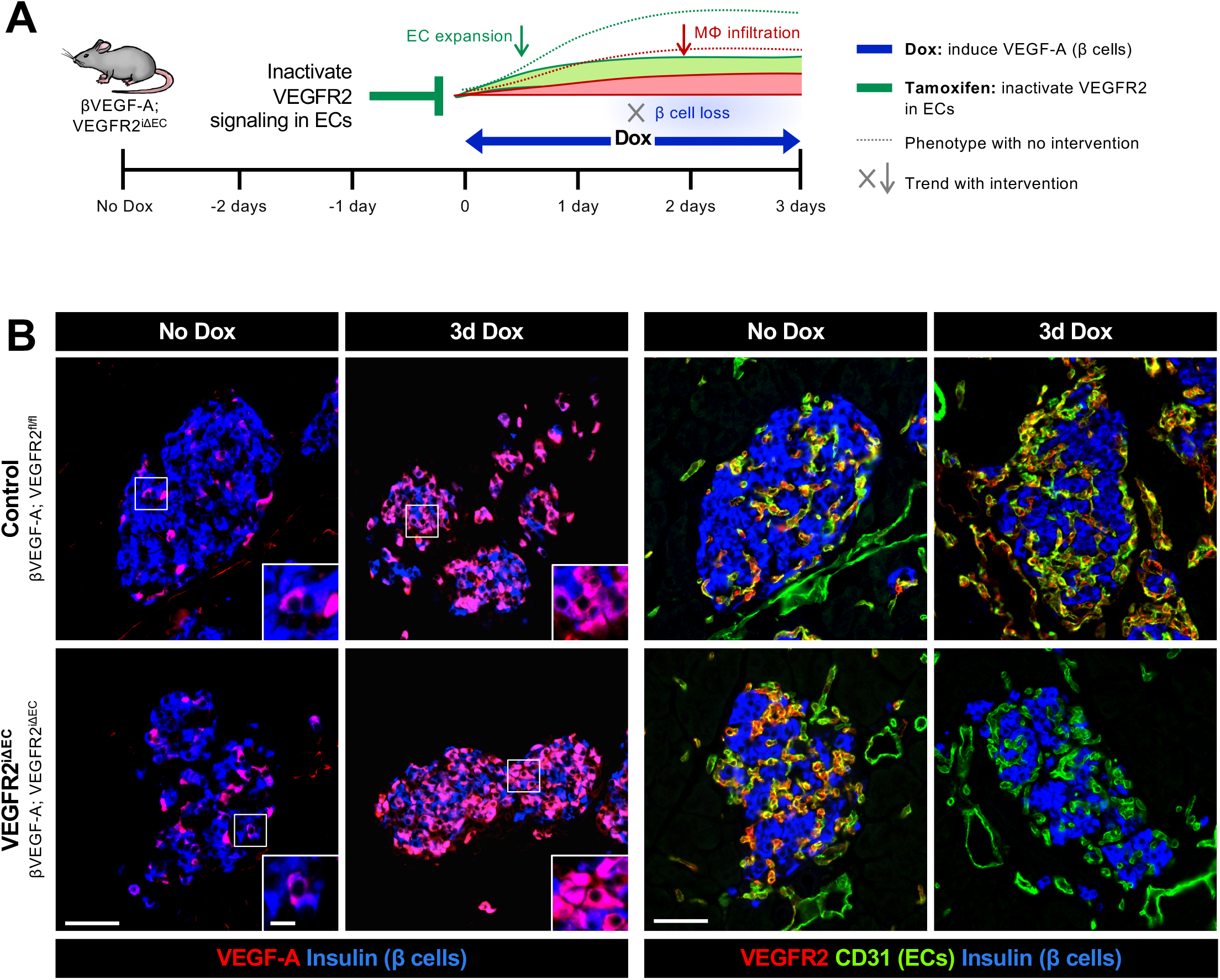
Effective VEGFR2 inactivation in endothelial cells prevents islet capillary expansion by acute elevation of VEGF-A in the islet microenvironment. **(A)** To inactivate VEGFR2 in endothelial cells (ECs), βVEGF-A; R2^iΔEC^ mice and βVEGF-A; R2^fl/fl^ controls were treated with Tamoxifen (Tm; 4mg s.c.) prior to VEGF-A induction. **(B)** Left panel: induction of VEGFA expression at 3d Dox compared to baseline (No Dox) in βVEGF-A; R2^iΔEC^ mice and controls; VEGF-A (red); β cells (Insulin^+^, blue). Boxes indicate regions enlarged in insets. Right panel: inactivation of VEGFR2 expression in ECs at baseline and 3d Dox; VEGFR2 (red), ECs (CD31^+^, green); β cells (Insulin^+^, blue). Scale bars, 50 μm; inset, 10 μm.

**Figure S5.**
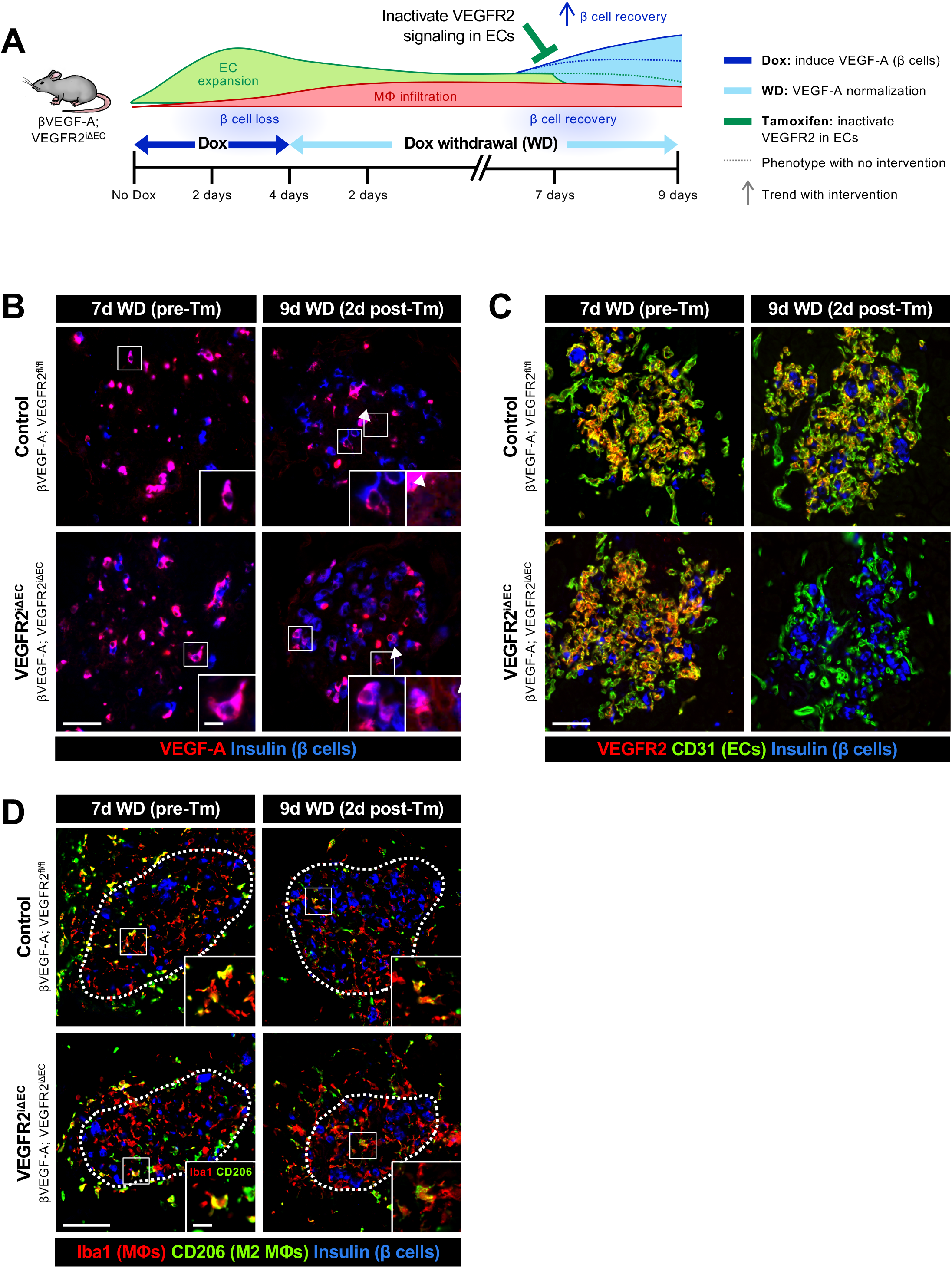
Effects of VEGF-A induction and VEGFR2 inactivation during β cell recovery. (A) To inactivate VEGFR2 in endothelial cells (ECs) during β cell recovery, control (βVEGF-A; VEGFR2^fl/fl^) and VEGFR2^iΔEC^ (βVEGF-A; VEGFR2^iΔEC^) mice received Tamoxifen (Tm; 4mg s.c.) after 7 days (d) of Dox withdrawal (WD). **(B)** Expression of VEGF-A in β cells and (**C**) VEGFR2 in ECs at 7d WD and 9d WD. Boxes show regions enlarged in insets; arrowheads point to extracellular VEGF-A staining. **(D)** Intra-islet M2-like macrophages (MΦs, CD206^+^ Iba1^+^) were present in both control and VEGFR2^iΔEC^ mice before and after Tm treatment. Approximate islet area is outlined; boxes show regions enlarged in insets with insulin channel (blue) removed. Scale bars in **(B–D)** are 50 μm; insets, 10 μm.

**Figure S6.**
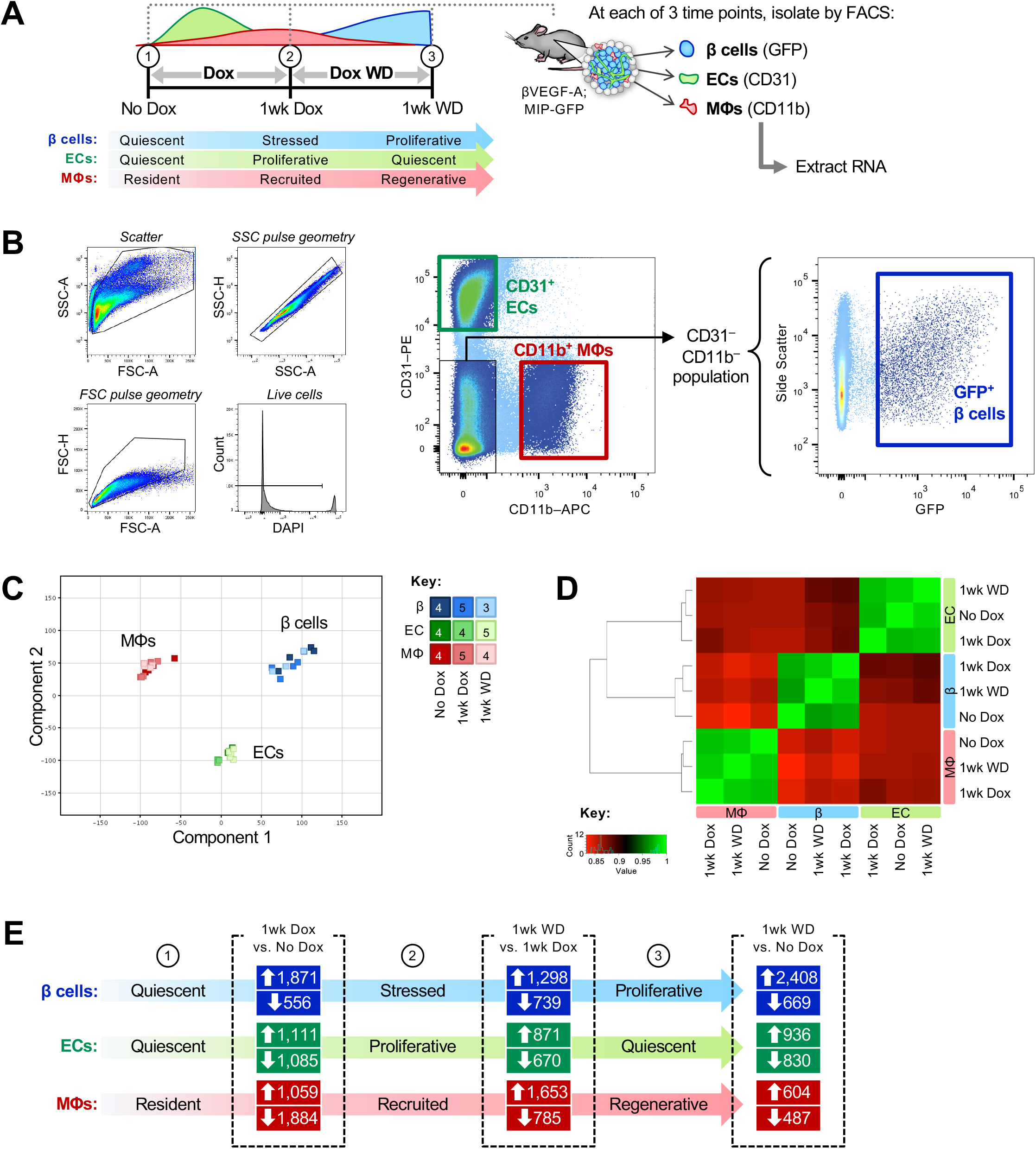
Transcriptome analysis demonstrates unique gene expression profiles in β cells, endothelial cells and macrophages purified from islets during the period of β cell loss and recovery. **(A–B)** Dispersed islet cells from βVEGF-A; MIP-GFP mice were immunolabeled with fluorophore-conjugated anti-CD31 and anti-CD11b antibodies to separate endothelial cell (EC) and macrophage (MΦ) populations, respectively. GFP^+^ β cells were sorted from the CD31^−^ CD11b^−^ population. Initial gating is shown in first four plots. **(C)**Principal component analysis (PCA) plot shows the clustering of samples from sorted β cells (blue), ECs (green), and MΦs (red) at No Dox, 1wk Dox, and 1wk WD; n=3-5 samples per time point as listed in Key. Islets from multiple mice were pooled to obtain adequate cells for each sample. **(D)** Pairwise correlation between all samples at all time points based on the Spearman correlation coefficient, which ranks and quantifies the degree of similarity between each pair of samples (perfect correlation=1; green). **(E)** Total number of genes for each cell type significantly up- or down-regulated (FC ≥2 or ≤−2) when comparing time points (dashed outlines): 1wk Dox vs. No Dox, 1wk WD vs. 1wk Dox, and 1wk WD vs. No Dox.

**Figure S7.**
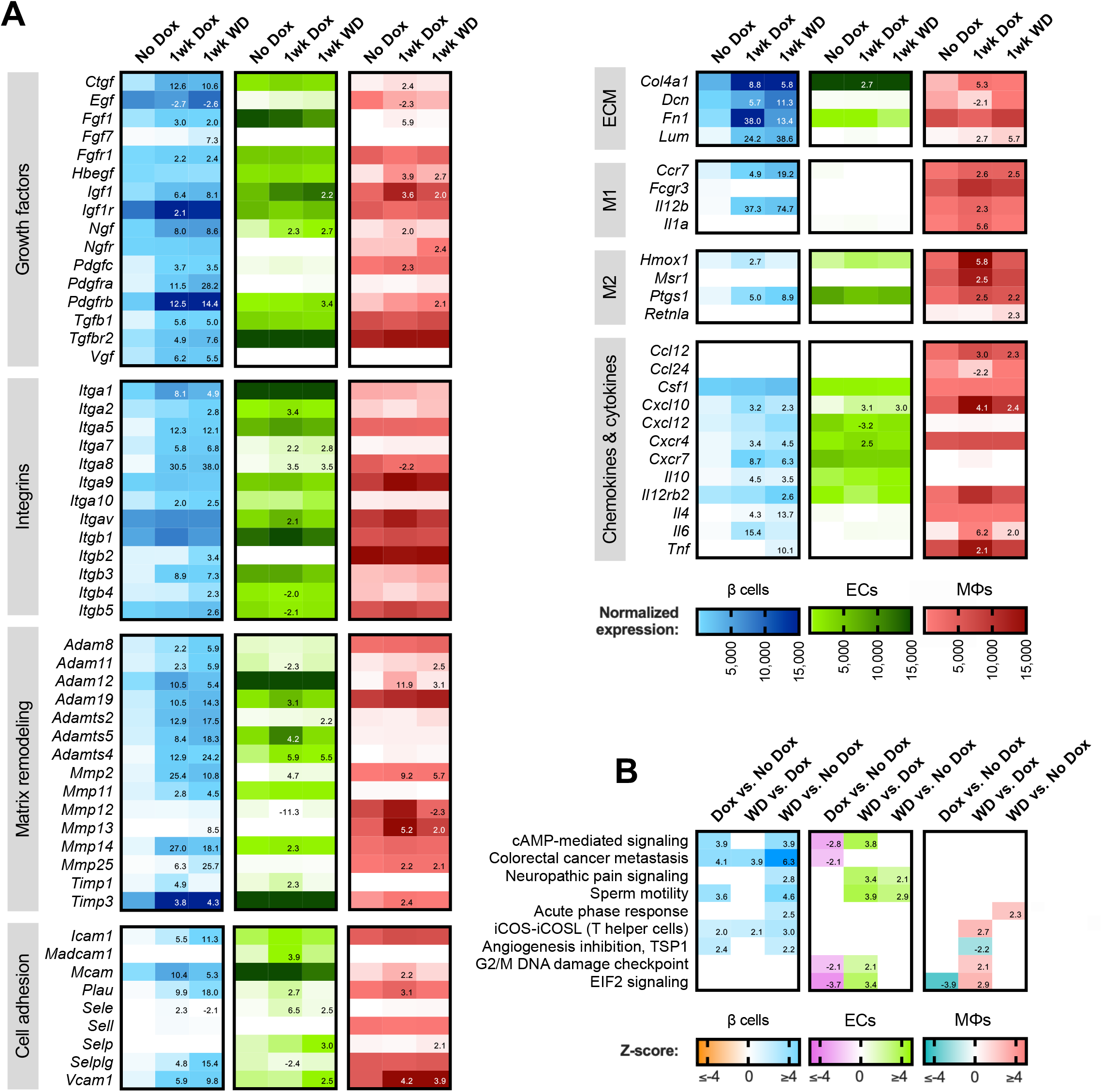
Temporal changes in gene expression profiles of islet cell subpopulations by RNA-sequencing. **(A)** Normalized expression of selected genes in isolated β cell (blue), EC (green), and MΦ (red) samples. ECM, extracellular matrix. Numbers listed on heat map in 1wk Dox and 1wk WD columns represent fold-change ≥2 or <−2 (p<0.05) as compared to No Dox. **(B)** Ingenuity Pathway Analysis (IPA) was applied to fold-change data (1wk Dox vs. No Dox, 1wk WD vs. 1wk Dox, and 1wk WD vs. No Dox) for β cell, EC, and MΦ populations. Pathways showing significant regulation (z-score ≥2 or ≤−2, p<0.05) in at least two cell types are shown; a full list is provided in **Table S1**. Z-scores of relevant comparisons are listed on heat map.

**Figure S8.**
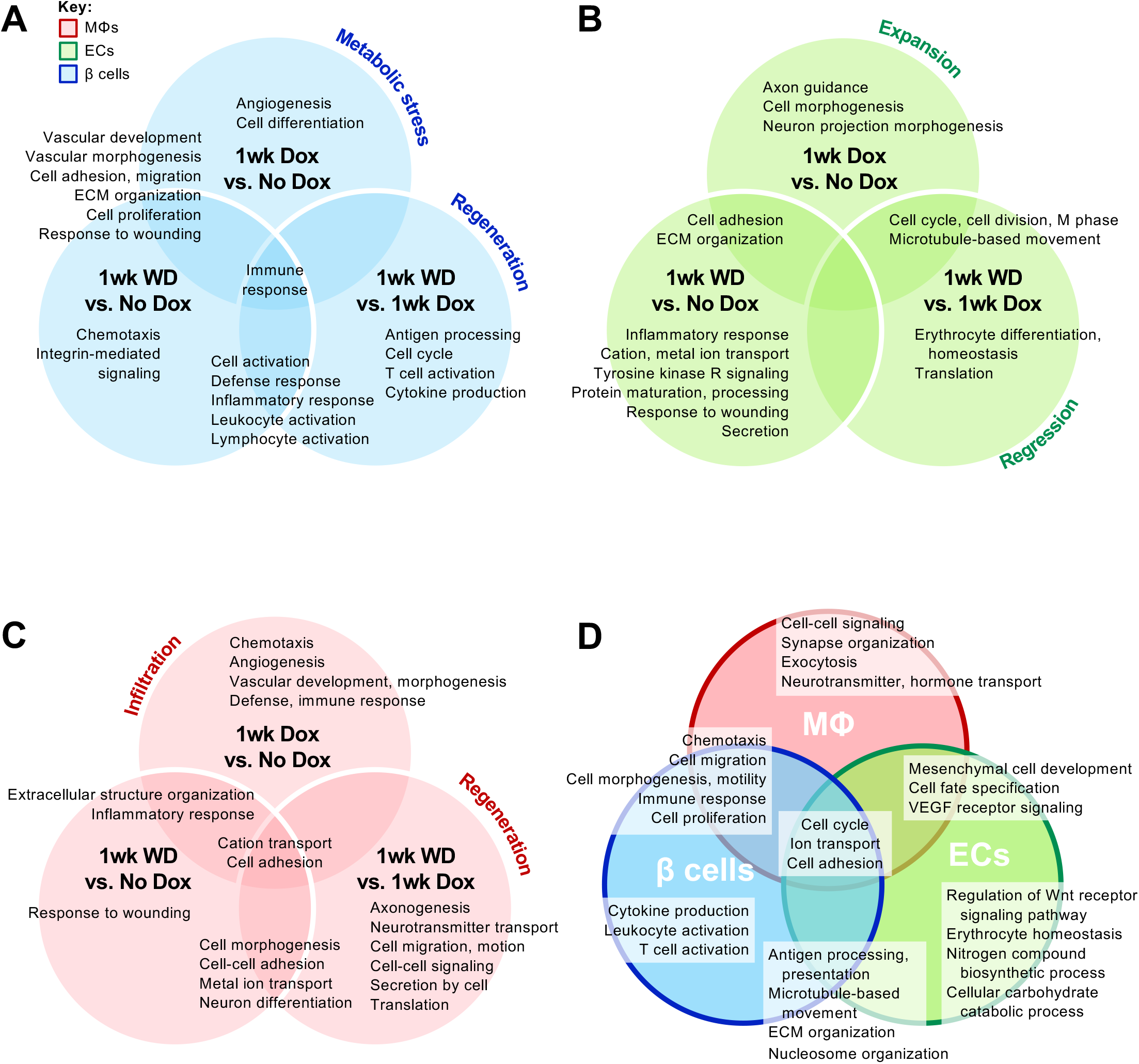
Biological processes enriched in islet macrophages, endothelial cells, and β cells of βVEGF-A mice during β cell loss and recovery. Venn diagrams showing profiles of β cells **(A)**, endothelial cells (ECs) **(B)**, and macrophages (MΦs) **(C)** as determined by Gene Ontology (GO) term analysis. Processes are organized by pairwise comparisons between time points (1wk Dox vs. No Dox, 1wk WD vs. 1wk Dox, and 1wk WD vs. No Dox), with location of text in Venn diagram accounting for processes that are represented in multiple time point comparisons. **(D)** Unique and shared processes of β cells, ECs, and MΦs during β cell recovery (1wk WD vs. 1wk Dox and/or 1wk WD vs. No Dox). All terms diagrammed in (**A–D**) appeared in top 20 hits (highest statistical significance) for the indicated population(s); those omitted due to space constraints are provided in **Table S2**.

**Table S1:**
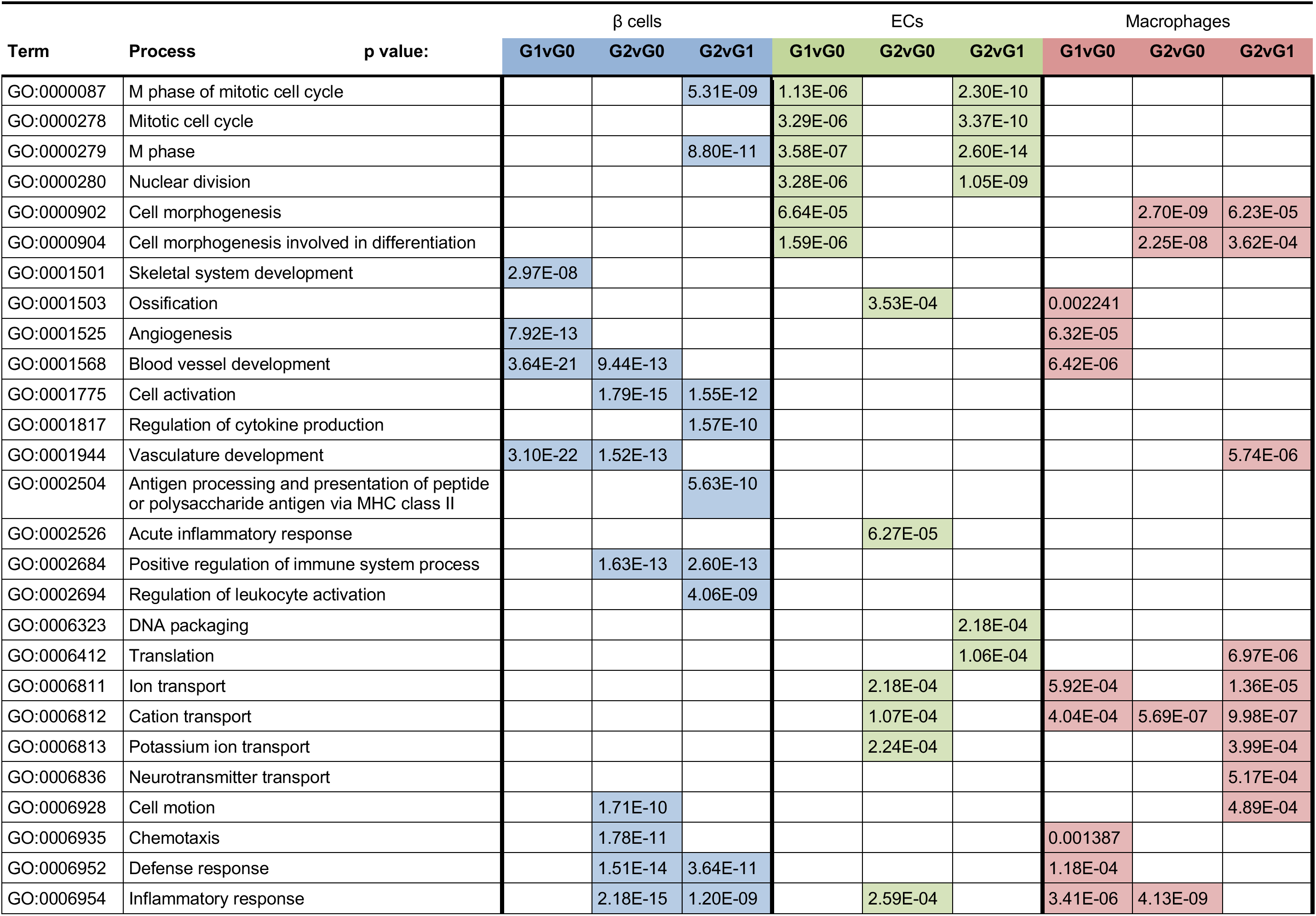
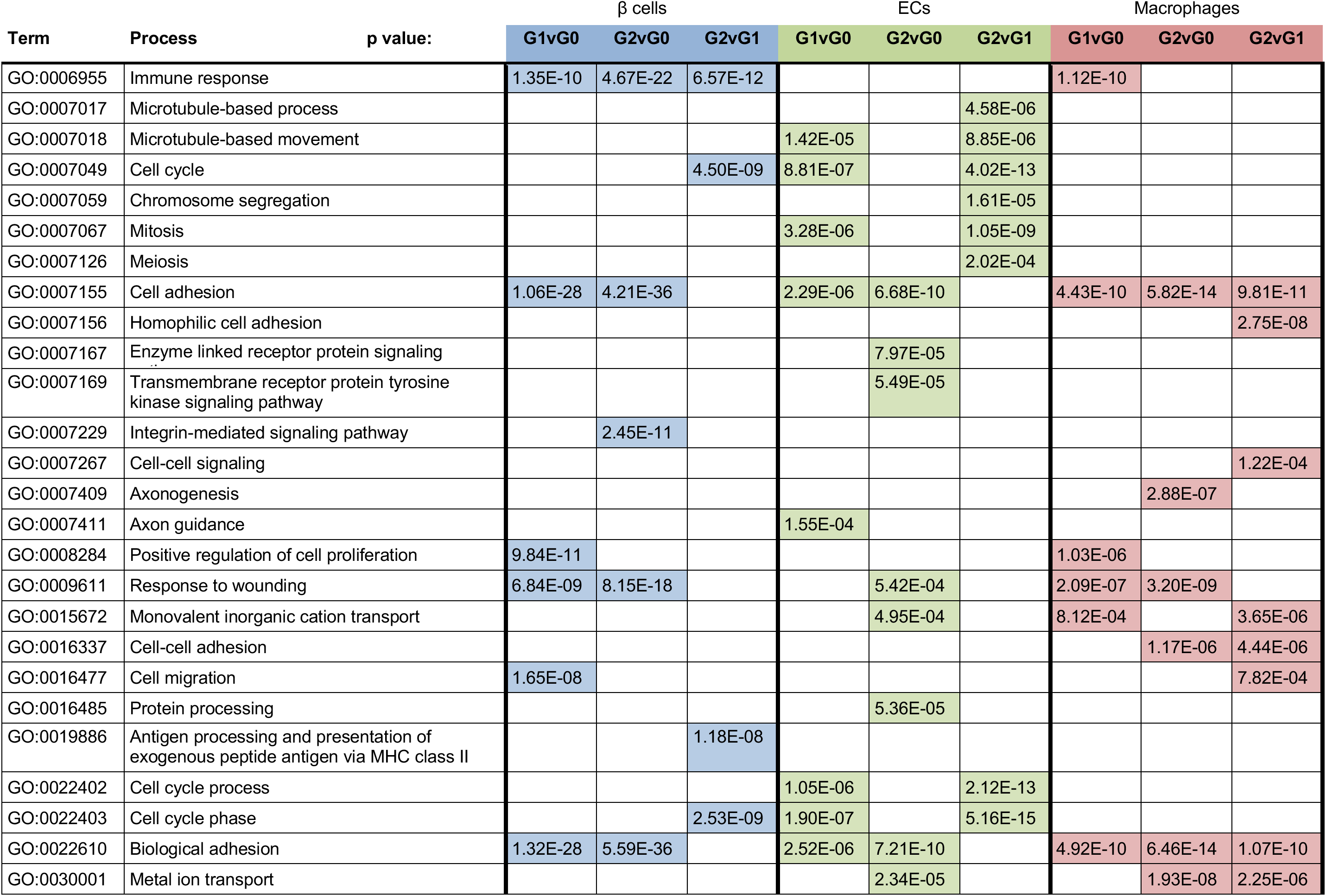
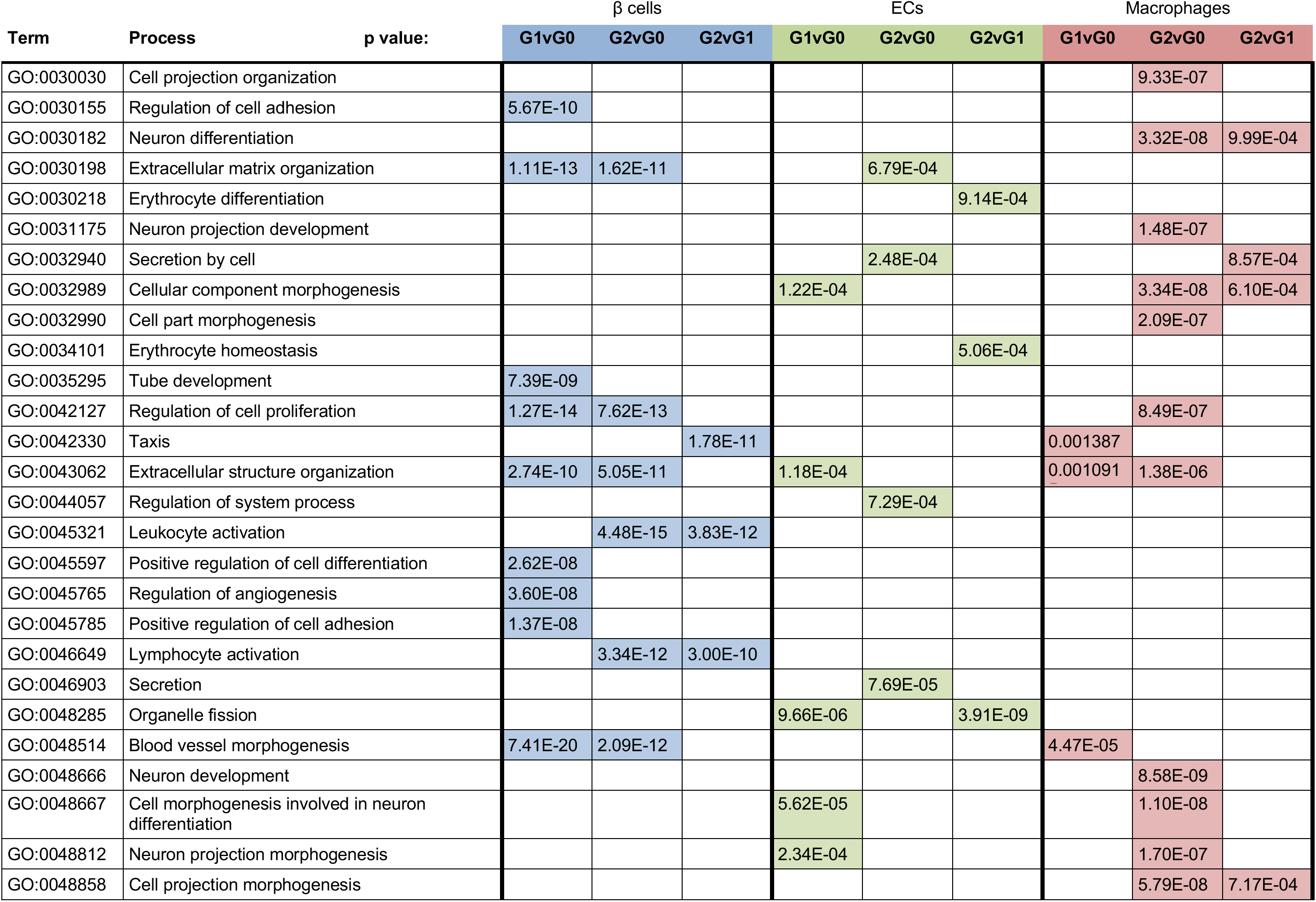
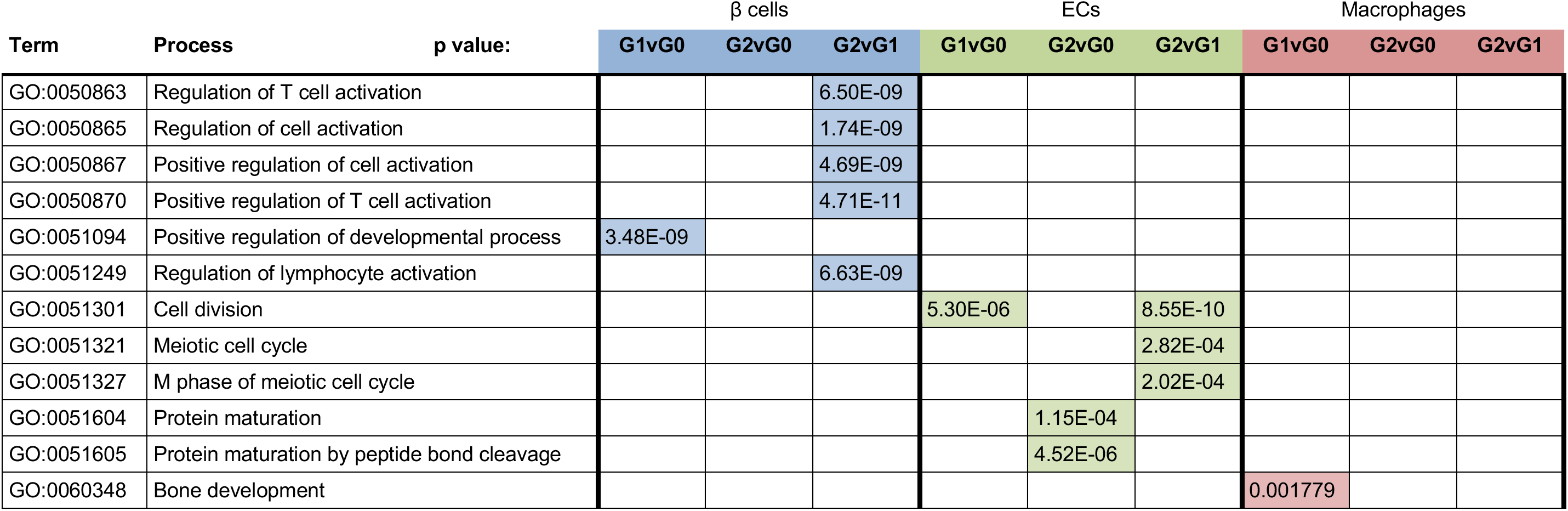
GO Terms for sorted cell populations

**Table S2:**
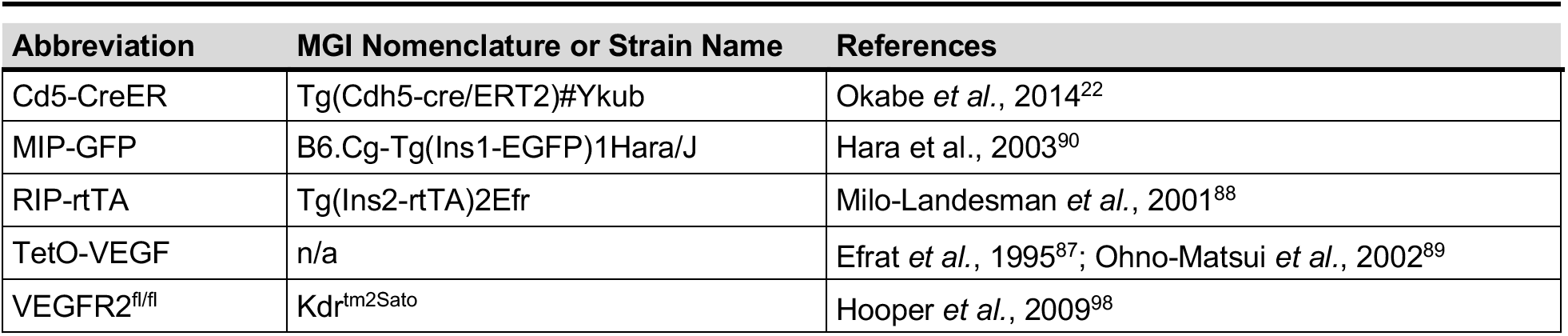
Mouse strains utilized in experimental models

| Abbreviation | MGI Nomenclature or Strain Name | References |
| --- | --- | --- |
| Cd5-CreER | Tg(Cdh5-cre/ERT2)#Ykub | Okabe *et al.*, 2014<sup>22</sup> |
| MIP-GFP | B6.Cg-Tg(Ins1-EGFP)1Hara/J | Hara *et al.*, 2003<sup>90</sup> |
| RIP-rtTA | Tg(Ins2-rtTA)2Efr | Milo-Landesman *et al.*, 2001<sup>88</sup> |
| TetO-VEGF | n/a | Efrat *et al.*, 1995<sup>87</sup>; Ohno-Matsui *et al.*, 2002<sup>89</sup> |
| VEGFR2<sup>fl/fl</sup> | Kdr<sup>tm2Sato</sup> | Hooper *et al.*, 2009<sup>98</sup> |

**Table S3:** Breeding scheme to generate βVEGF-A; VEGFR2^iΔEC^ mice

| Cross | Male breeder | Female breeder | Desired Offspring | Frequency |
| --- | --- | --- | --- | --- |
| A1 | Cd5-CreER<sup>tg/+</sup> | VEGFR2<sup>fl/fl</sup> | Cd5-CreER<sup>tg/+</sup>; VEGFR2<sup>fl/+</sup> | 50% |
| A2 | Cd5-CreER<sup>tg/+</sup>; VEGFR2<sup>fl/+</sup> | VEGFR2<sup>fl/fl</sup> | Cd5-CreER<sup>tg/+</sup>; VEGFR2<sup>fl/fl</sup> | 25% |
| A3 | Cd5-CreER<sup>tg/+</sup>; VEGFR2<sup>fl/fl</sup> | RIP-rtTA<sup>tg/tg</sup> | Cd5-CreER<sup>tg/+</sup>; VEGFR2<sup>fl/+</sup>; RIP-rtTA<sup>tg/+</sup> | 50% |
| A4 | Cd5-CreER<sup>tg/+</sup>; VEGFR2<sup>fl/+</sup>; RIP-rtTA<sup>tg/+</sup> | VEGFR2<sup>fl/fl</sup> | Cd5-CreER<sup>tg/+</sup>; VEGFR2<sup>fl/fl</sup>; RIP-rtTA<sup>tg/+</sup> | 12.5% |
| B1 | VEGFR2<sup>fl/fl</sup> | TetO-VEGFA<sup>tg/tg</sup> | TetO-VEGFA<sup>tg/+</sup>; VEGFR2<sup>fl/+</sup> | 100% |
| B2 | TetO-VEGFA<sup>tg/+</sup>; VEGFR2<sup>fl/+</sup> | TetO-VEGFA<sup>tg/tg</sup> | TetO-VEGFA<sup>tg/tg</sup>; VEGFR2<sup>fl/+</sup> | 50% |
| B3 | TetO-VEGFA<sup>tg/+</sup>; VEGFR2<sup>fl/+</sup> | TetO-VEGFA<sup>tg/tg</sup>; VEGFR2<sup>fl/+</sup> | TetO-VEGFA<sup>tg/tg</sup>; VEGFR2<sup>fl/fl</sup> | 25% |
| C1 | Cd5-CreER<sup>tg/+</sup>; VEGFR2<sup>fl/fl</sup>; RIP-rtTA<sup>tg/+</sup> | TetO-VEGFA<sup>tg/tg</sup>; VEGFR2<sup>fl/fl</sup> | TetO-VEGFA<sup>tg/+</sup>; Cd5-CreER<sup>tg/+</sup>; VEGFR2<sup>fl/fl</sup>; RIP-rtTA<sup>tg/+</sup> | 25% |

**Table S4:**
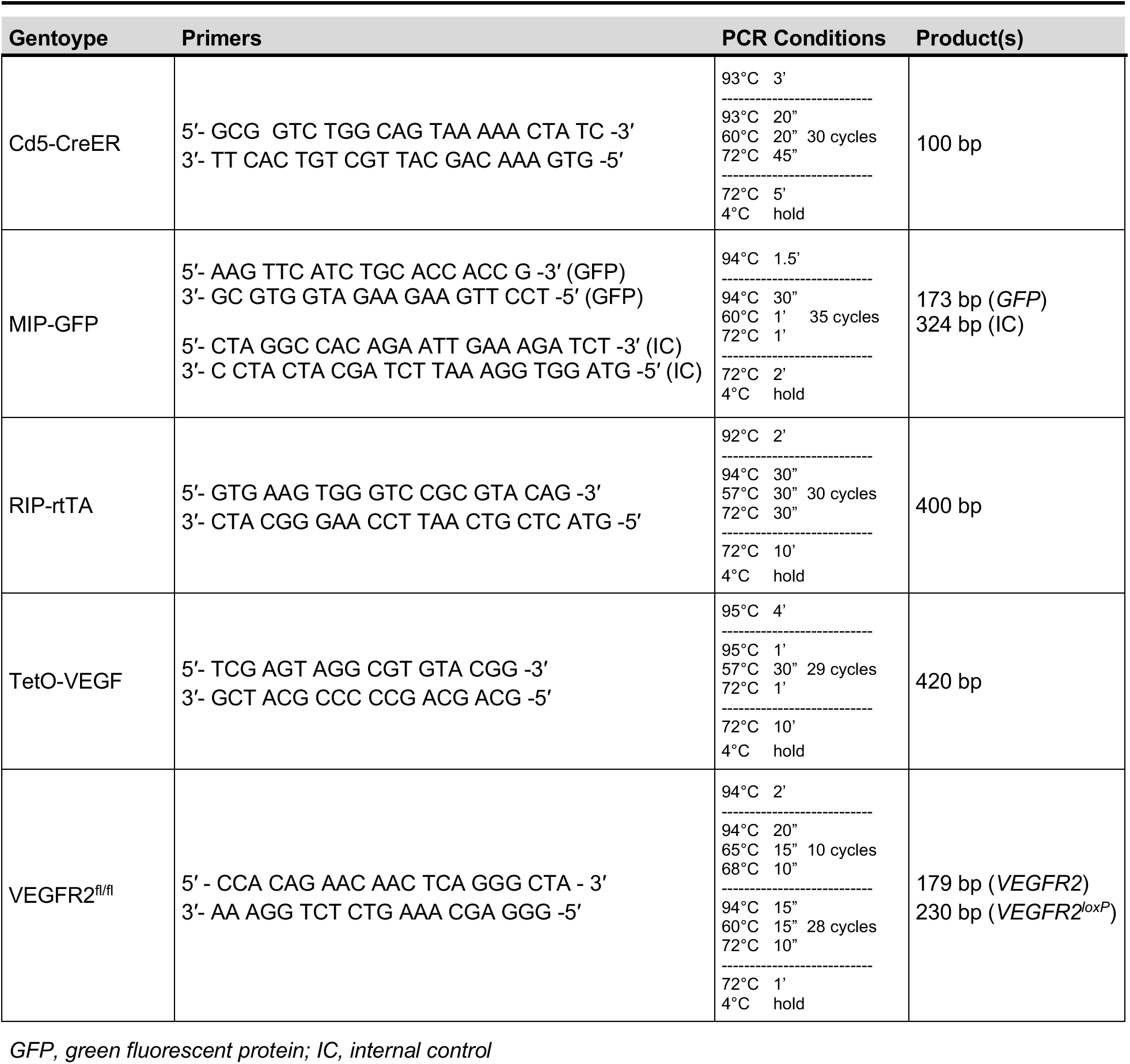
PCR primers and conditions for genotyping

**Table S5:** Antibodies used for immunohistochemistry and flow cytometry

| Antigen/Conjugate | Species | Source | Catalog # | Application | Dilution |
| --- | --- | --- | --- | --- | --- |
| Caveolin-1 | Rabbit | Abcam | ab2910 | IHC (1°) | 1:2000 |
| CD11b – APC | Rat | BD Pharmingen | 561690 | FC | 1:100,<br>1:500 |
| CD206 (Mrc1) | Rat | BioLegend | 141701 | IHC (1°) | 1:500 |
| CD31 | Rat | BD Pharmingen | 553370 | IHC (1°) | 1:100 |
| CD31 – PE | Rat | BD Pharmingen | 561073 | FC | 1:500 |
| CD45 – PE | Rat | BD Pharmingen | 561087 | FC | 1:500 |
| Col-IV | Rabbit | Abcam | ab6586 | IHC (1°) | 1:1000 |
| Goat IgG – Cy3 | Donkey | Jackson Immunoresearch | 705-165-147 | IHC (2°) | 1:500 |
| Guinea pig IgG – Cy2 | Donkey | Jackson Immunoresearch | 706-225-148 | IHC (2°) | 1:500 |
| Guinea pig IgG – Cy5 | Donkey | Jackson Immunoresearch | 706-175-148 | IHC (2°) | 1:200 |
| Iba1 | Rabbit | Wako | 019-19741 | IHC (1°) | 1:500 |
| Iba1 | Rabbit | Abcam | ab221790 | IHC (1°) | 1:1000 |
| Insulin | Guinea pig | Dako | A0564 | IHC (1°) | 1:1000 |
| Insulin | Guinea pig | Cell Marque | 273A-15 | IHC (1°) | 1:500 |
| Ki67 | Rabbit | Abcam | ab15580 | IHC (1°) | 1:5000 |
| Ly6G-FITC | Rat | BD Pharmingen | 551460 | FC | 1:500 |
| pERK | Rabbit | Cell Signaling | 4370 | IHC (1°) | 1:200 |
| Rabbit IgG – Cy2 | Donkey | Jackson Immunoresearch | 711-225-152 | IHC (2°) | 1:500 |
| Rabbit IgG – Cy3 | Donkey | Jackson Immunoresearch | 711-165-152 | IHC (2°) | 1:500 |
| Rat IgG – Cy2 | Donkey | Jackson Immunoresearch | 712-225-153 | IHC (2°) | 1:500 |
| VEGF-A | Goat | R&D Systems | AF564 | IHC (1°) | 1:200 |
| VEGFR2 | Goat | R&D Systems | AF644 | IHC (1°) | 1:2000 |

1°, primary antibody; 2°, secondary antibody; FC, flow cytometry; IHC, immunohistochemistry

